# Global loss of cellular m^6^A RNA methylation following infection with different SARS-CoV-2 variants

**DOI:** 10.1101/2022.12.08.519593

**Authors:** Roshan Vaid, Akram Mendez, Ketan Thombare, Rebeca Burgos-Panadero, Rémy Robinot, Barbara F Fonseca, Nikhil R Gandasi, Johan Ringlander, Mohammad Hassan Baig, Jae-June Dong, Jae Yong Cho, Björn Reinius, Lisa A Chakrabarti, Kristina Nystrom, Tanmoy Mondal

**Author notes:** Equal contribution (Roshan Vaid, Akram Mendez). Corresponding author,. Department of Clinical Chemistry and Transfusion Medicine, Bruna Stråket 16, Sahlgrenska University Hospital, Gothenburg University, Gothenburg, 41345, Sweden.

## Abstract

Host-viral interactions during SARS-CoV-2 infection are needed to understand COVID-19 pathogenesis and may help to guide the design of novel antiviral therapeutics. *N*^6^-methyladenosine modification (m^6^A), one of the most abundant cellular RNA modifications, regulates key processes in RNA metabolism during a stress response. Gene expression profiles observed post-infection with different SARS-CoV-2 variants show changes in the expression of genes related to RNA catabolism, including m^6^A readers and erasers. We found that infection with SARS-CoV-2 variants caused a loss of m^6^A in cellular RNAs, whereas m^6^A was detected abundantly in viral RNA. METTL3, the m^6^A methyltransferase, showed an unusual cytoplasmic localization post-infection. The B.1.351 variant had a less pronounced effect on METTL3 localization and loss of m^6^A than the B.1 and B.1.1.7 variants. We also observed a loss of m^6^A upon SARS-CoV-2 infection in air/liquid interface cultures of human airway epithelia, confirming that m^6^A loss is characteristic of SARS-CoV-2 infected cells. Further, transcripts with m^6^A modification were preferentially down-regulated post-infection. Inhibition of the export protein XPO1 resulted in the restoration of METTL3 localization, recovery of m^6^A on cellular RNA, and increased mRNA expression. Stress granule formation, which was compromised by SARS-CoV-2 infection, was restored by XPO1 inhibition and accompanied by a reduced viral infection *in vitro*. Together, our study elucidates how SARS-CoV-2 inhibits the stress response and perturbs cellular gene expression in an m^6^A-dependent manner.

## INTRODUCTION

The ongoing pandemic of COVID-19 disease is caused by the severe acute respiratory syndrome coronavirus 2 (SARS-CoV-2). Since 2019, the virus has spread over the world and novel variants have emerged in succession, associated with changes in transmissibility and disease severity capacity (Tao et al. 2021). The SARS-CoV-2 genome consists of a single-stranded positive genomic RNA of approximately 30,000 nucleotides (Wu et al. 2020). Whereas different vaccines have been developed to mitigate SARS-CoV-2 spreading, it is still necessary to gain knowledge on the basic mechanisms underlying SARS-CoV-2 infection and host response to assist the development of improved therapeutic options.

*N*^6^-methyladenosine modification (m^6^A) is a prevalent internal RNA modification (Baquero-Perez et al. 2021). It is involved in the regulation of a broad range of biological processes including cell differentiation (Geula et al. 2015), mRNA stability, translation, liquid-phase separation, and stress granule formation, among others (Zaccara et al. 2019). The m^6^A modification is deposited by a protein complex consisting of m^6^A writers. The writing of m^6^A on mRNA is mediated by a methyltransferase complex comprising the core catalytic subunits METTL3 and METTL14, and the adapter subunits WTAP, VIRMA, HAKAI, and RBM15/B (Patil et al. 2016; Xiao et al. 2016; Bawankar et al. 2021; Yue et al. 2018; Ping et al. 2014). The m^6^A-modified RNAs are directly recognized by cytoplasmic YTH domain-containing proteins, namely YTHDF1-3, which regulate mRNA stability and stress granule formation. In addition, the nuclear m^6^A reader proteins YTHDC1 and hnRNPA2B1 play critical roles in splicing and nuclear export (Xiao et al. 2016; Alarcón et al. 2015). Finally, m^6^A modifications can be erased by the dioxygenases FTO and ALKBH5, which specifically demethylate m^6^A RNA and thus control the reversibility of m^6^A modifications (Wei et al. 2022; Zhang et al. 2017)

The m^6^A RNA modification has been identified in several viral genomic RNA, being first described for DNA viruses such as the simian virus 40 (SV40), herpes simplex virus, and adenovirus 2, and also in retroviruses like Rous sarcoma virus, and human immunodeficiency virus 1 (HIV-1) (Lu et al. 2018). Moreover, a role for m^6^A modification has been described for RNA viruses such as Dengue, West Nile fever, Yellow fever, Zika, and Hepatitis C viruses, where it was involved in the suppression of viral gene expression and replication (Kim and Siddiqui 2021; Lichinchi et al. 2016). The SARS-CoV-2 genome is m^6^A-modified by host proteins, and the m^6^A modification is important for promoting viral replication and for limiting the host immune response (Liu et al. 2021; Li et al. 2021; Zhang et al. 2021b). Depletion of the cytoplasmic m^6^A readers, the YTHDF proteins, as well as the writer METTL3, suppress SARS-CoV-2 (and HCoV-OC43) replication. METTL3 inhibition mediated by a small molecule inhibitor induces viral RNA synthesis suppression, suggesting that m^6^A modification is needed for optimal viral RNA expression (Burgess et al. 2021).

Though recent studies reported the presence of m^6^A in the SARS-CoV-2 RNA genome (Li et al. 2021; Zhang et al. 2021b; Kim et al. 2020; Burgess et al. 2021; Liu et al. 2021), it has not been explored in detail how the host’s m^6^A mRNA profile is affected during infection. To address this question, we studied the effect of SARS-CoV-2 infection in Vero cells and air/liquid interface (ALI) cultures of human airway epithelia, which are highly permissive to SARS-CoV-2 infection (Wei et al. 2021b; Robinot et al. 2021).

## RESULTS

### Infection by different SARS-CoV-2 variants leads to deregulation of RNA catabolism related genes

To identify variant-specific differences in gene expression perturbations, we infected Vero cells with three SARS-CoV-2 variants: B.1, B.1.1.7, and B.1.351. At 48 h post-infection, spike positivity was detected in 94-95% of infected cells for all the variants (**Supplemental Fig. S1A**). We isolated total RNA 48-h post-infection and evaluated gene expression changes using RNA-sequencing (RNA-seq). We first checked viral RNA levels post-infection and observed that RNA-seq reads mapping to the viral genome were comparable for the three SARS-CoV-2 variants (**Supplemental Fig. S1B**). The expression levels analyzed per viral gene were also similar for the three variants (**Supplemental Fig. S1C**). Viral reads contributed 1.2 to 2.8% of the total reads recovered from infected cells (adding reads from the viral and host genomes) (**Supplemental Fig. S1D),** which is consistent with an earlier report (Blanco-Melo et al. 2020). Next, we performed a differential gene expression analysis to identify genes with altered expression post-infection in Vero cells (**Fig. 1A; Supplemental Data S1**). We identified a considerable number of up- (998) and down-regulated (950) genes that were modulated across the three variants during infection (**Fig. 1B).** We then performed a principal component analysis (PCA) and a regression analysis to evaluate similarities in gene expression patterns across the three variants. We observed variant-specific patterns in gene expression, with gene expression patterns being more similar between the B.1.1.7 and B.1.351 variants (**Supplemental Fig. S1E, F**). Gene expression changes post-infection with three SARS-CoV-2 variants overlapped with several publicly available datasets deposited since the outbreak of COVID-19 (**Supplemental Fig. S1G**). We also compared the gene expression dataset post-SARS-CoV-2 infection in three different variants with one of the publicly available data sets (Riva et al. 2020), where differential gene expression was analyzed 24 h post-SARS-COV-2 infection in the Vero cells, and we observed moderate but significant positive correlation between our data and publicly available data (**Supplemental Fig. S1H**). Pathway analysis revealed that altered genes were enriched in pathways associated with protein localization to the membrane, RNA-catabolic processes, and cilium assembly (**Fig. 1C; Supplemental Data S2**). This was evident in the pathway analysis performed on up- and down-regulated genes separately post-viral infection. Up-regulated genes were enriched for pathways related to several RNA catabolic processes, especially in B.1 and B.1.1.7 variants whereas down-regulated genes were enriched with cilium assembly and organization (**Supplemental Fig. S1**I; **Supplemental Data S2**). We then chose to systematically look at the genes involved in RNA catabolic processes that were frequently deregulated across variants during SARS-CoV-2 infection. To do so, we selected RNA catabolism-related pathways which were enriched in deregulated genes in at least two variants and visualized the interactions of those genes using network analysis (**Fig. 1D**). We observed that the mean connectivity of nodes in the RNA catabolism pathway-related network was much higher compared to random networks (**Supplemental Fig. S1J**). In particular, the network analysis suggested widespread interactions (as detected by connectivity degree) of m^6^A-related genes with different RNA catabolic processes (**Supplemental Fig. S1K**). We observed that m^6^A-related genes were frequently deregulated during SARS-CoV-2 infection in Vero cells (**Fig. 1E**), though we did not observe any significant change in the expression of the main m^6^A writers *METTL3* and *METTL14*. We validated the differential expression of m^6^A eraser *FTO*, which was down-regulated, and the up-regulation of the m^6^A-related genes *SPEN* and *YTHDF1* (Dossin et al. 2020), which were observed after infection with the three variants. Similar changes were observed after infection with the B.1.617.2 (Delta) variant, which was available during the final compilation of the data (**Fig. 1F**). Though we observed variant-specific gene expression changes, our data suggest widespread deregulation of genes associated with RNA catabolic processes and m^6^A modification-related pathways for all the SARS-CoV-2 variants tested.

**Fig. 1.**
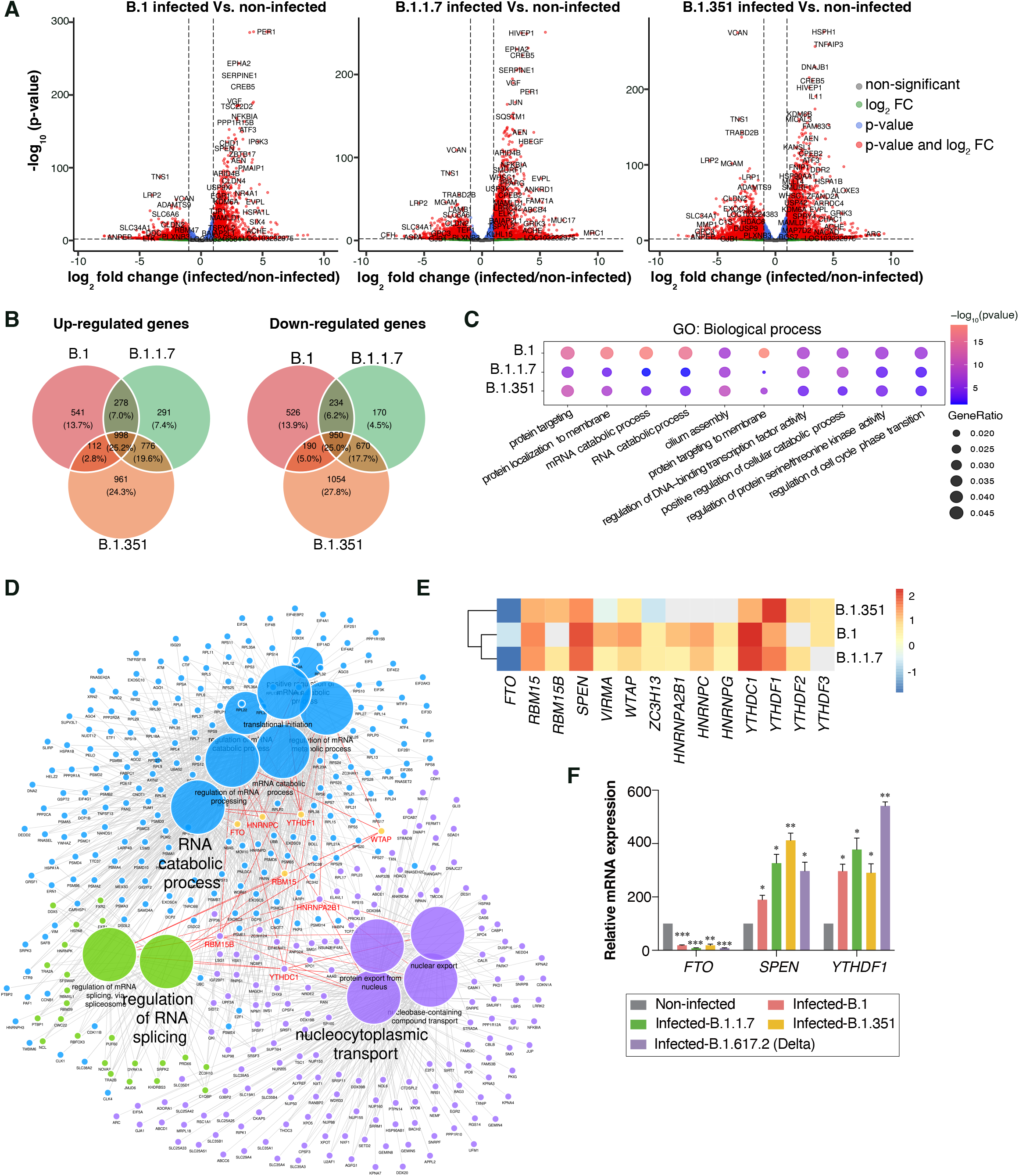
Gene expression changes post-infection with different variants of SARS-CoV-2. (A) Volcano plots showing gene expression changes and significance (-log_10_ scale) in SARS-CoV-2 infected versus non-infected cells. Dashed lines indicate the significance threshold (adjusted p-value < 0.01) and log_2_ fold change threshold (abs log_2_ fold change > 1). Differentially expressed genes are highlighted in red, and labels indicate common differentially expressed genes across all conditions. (B) Venn diagram comparison of up- and down-regulated genes across variants. (C) Top enriched GO term biological processes after infection. The size of the dots represents the enrichment of genes with a GO term, colored according to their significance level. (D) ClueGO clustering and visualization of common terms associated with m^6^A-related genes across any pair of SARS-CoV-2 variants. m^6^A genes that are deregulated across variants are highlighted in red. (E) Heat map of m^6^A-related genes, with grey boxes indicating non-differentially expressed genes whereas colors indicate log_2_ fold change values of significantly deregulated genes in at least two variants. (F) Relative mRNA expression of m^6^A-related genes in non-infected Vero cells and Vero cells infected with B.1, B.1.1.7, B.1.351, and B.1.617.2 (Delta) variants of SARS-CoV-2. *TBP* and *POL2RG* were used to normalize the qPCR data. Data are shown as Mean ± SD of three replicates (n=3). Statistics: two-tailed paired *t*-test, *: p < 0.05, **: p < 0.01, ***: p < 0.001.

### Variant-specific changes in cellular RNA m^6^A level after viral infection

A change in m^6^A-related genes during viral infection prompted us to check the effect of infection on cellular m^6^A levels. To this end, isolated RNA from non-infected Vero cells and cells infected with three SARS-CoV-2 variants (B.1, B.1.1.7, and B.1.351) was supplemented with spike-in bacterial RNA as a control (spike-in) for m^6^A RNA Immunoprecipitation followed by Sequencing (m^6^A RIP-seq) (**Fig. 2A**). We observed a loss of m^6^A peaks in the cellular RNA post-infection with different SARS-CoV-2 variants, with the loss of m^6^A peaks being more marked in B.1 and B.1.1.7 infections compared to B.1.351 infection (**Supplemental Fig. S2A; Supplemental Data S3)**. The global distribution of m^6^A peak density across transcripts in non-infected cells was similar to that reported previously, with strong enrichments near the start and stop codons (Meyer et al. 2012). Spike-in normalized relative m^6^A peak density showed m^6^A loss across the whole transcript length following SARS-CoV-2 infection in Vero cells, with more pronounced effects in B.1 and B.1.1.7 compared to B.1.351 infection (**Fig. 2B**). This global loss of m^6^A was also evident by the reduced number of genes with m^6^A peaks following SARS-CoV-2 infection (**Supplemental Fig. S2B**). A decrease in the spike-in normalized m^6^A signal was visible over the commonly lost m^6^A peaks (**Supplemental Fig. S2C)** and also at the level of individual transcripts (**Fig. 2C)** after infection with three SARS-CoV-2 variants. Identified m^6^A-peaks in non-infected cells were enriched with the known “GGAC” motif, whereas the dominant motif was more degenerate in infected cells (**Fig. 2D**). Although the number of m^6^A peaks was drastically reduced, we also detected new m^6^A peaks that were gained over many transcripts after infection with all three variants. The number of gained m^6^A peaks was highest for the B.1.351 variant (**Supplemental Fig. S2B; Fig. 2E**). When we investigated the genomic location of the lost and gained peaks during infection, we observed that gained peaks were found in different genomic locations with major gained peaks located at intergenic and TSS regions (**Supplemental Fig. S2E**). The most enriched motifs (despite less significant p-values) in the gained m^6^A peaks were different from the canonical “GGAC” motif (**Supplemental Fig. S2D**). We also checked by scatter plots the relation between expression (log_2_ fold changes: infected vs non-infected) and m^6^A enrichment (infected vs non-infected) across the transcripts and found that change in expression cannot simply explain alteration in m^6^A enrichment (**Supplemental Fig. S2F**). This was also evident in the input RNA signal, which was mostly unaffected over the common lost peaks in infected cells even though the m^6^A signal was reduced over those peaks (**Supplemental Fig. S2C)**, suggesting the involvement of additional mechanisms which drive m^6^A loss during SARS-CoV-2 infection.

**Fig. 2.**
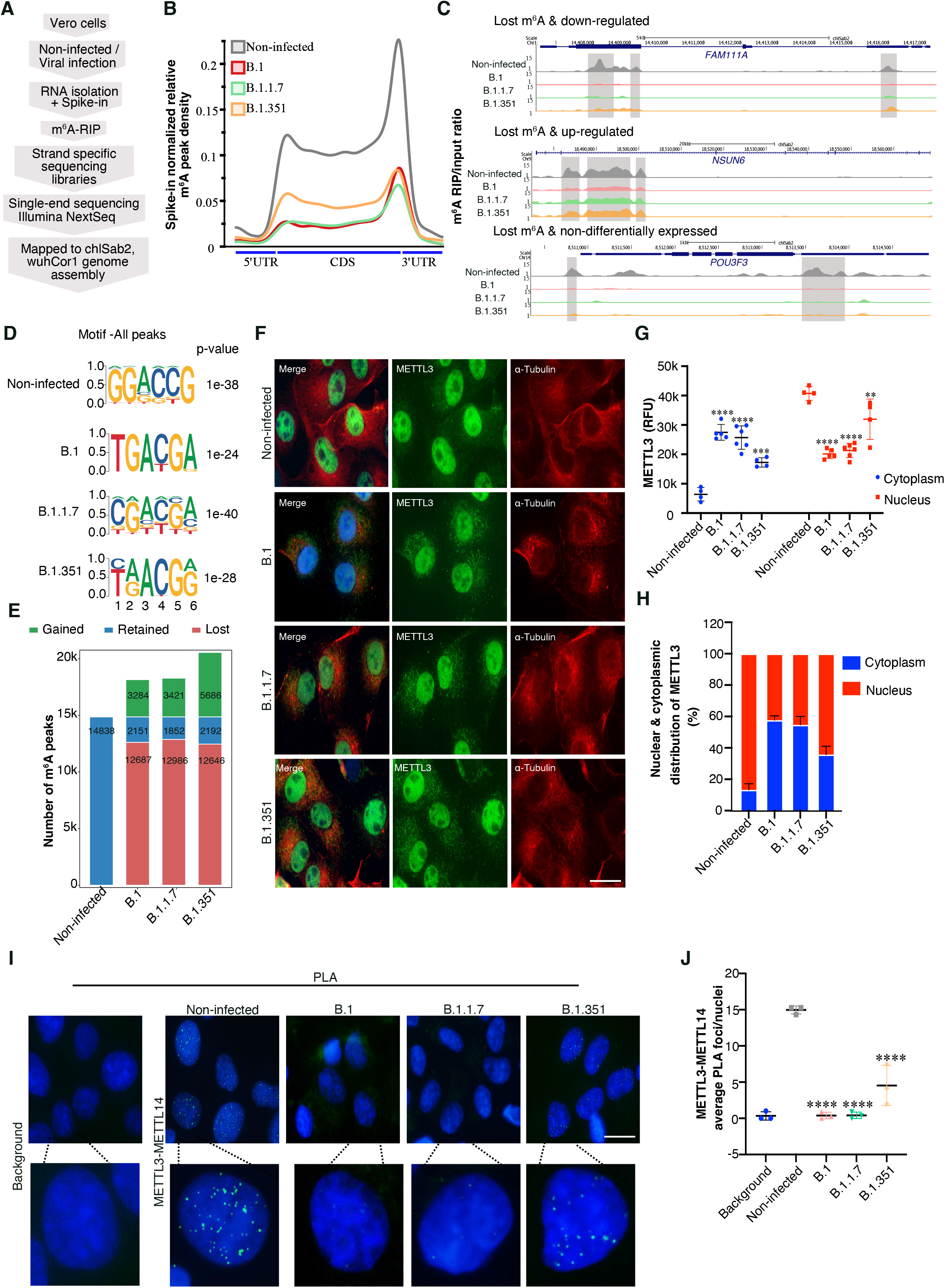
Variant-specific changes in cellular RNA m^6^A level after viral infection. (A) Flowchart describing the m^6^A RIP-seq protocol. (B) Metagene analysis showing spike-in normalized relative m^6^A peak density at host genes in non-infected and SARS-CoV-2 infected Vero cells (variants as specified by colors). (C) Genome browser screenshots showing spike-in normalized m^6^A (IP/input ratio) tracks for three different genes in non-infected and SARS-CoV-2 infected (variants as specified) Vero cells. m^6^A peak regions identified using the MACS peak caller software in non-infected cells are highlighted using gray boxes. (D) Identified motifs from *de novo* motif analysis of peaks in non-infected and infected cells. (E) The total number of peaks classified as retained, gained, or lost, post-infection with the SARS-CoV-2 variant are indicated and compared to the non-infected condition. (F-H) METTL3 localization: (F) METTL3 and α-Tubulin co-immunostaining in Vero cells that were either non-infected or infected with SARS-CoV-2 variants (as denoted). Scale bar is 20 μm. (G) METTL3 intensities (in Relative Fluorescence Units: RFU) derived from the immunostainings performed in (F). METTL3 RFU in the nucleus (red) and cytoplasm (blue) were estimated using ImageJ, using DAPI marking the nucleus and α-Tubulin staining the cytoplasm as references. Data are shown as Mean ± SD. Data presented from multiple experiments with the total number of cells counted, n=<70. One-way ANOVA test was performed, **: p < 0.01, ***: p < 0.001, ****: p < 0.0001. (H) Distribution of METTL3 in the nucleus and cytoplasm calculated from the percentage of RFU intensities measured as described in (G). Data are shown as Mean ± SD. (I-J) METTL3-METTL14 interaction: (I) Proximity ligation assay (PLA) in non-infected and SARS-CoV-2 infected Vero cells (variants as indicated) depicting METTL3 and METTL14 PLA foci in the nucleus (marked by DAPI). The background control shows PLA with only the METTL3 antibody. Scale bar is 20 μm. (J) Quantification of METTL3-METTL14 PLA foci as detected in (I). The number of PLA foci/nuclei are shown as Mean ± SD. Data presented from multiple experiments with the total number of cells counted, n=<100. One-way ANOVA test was performed, ****: p < 0.0001.

The global loss of m^6^A peaks during infection suggests an altered function of the key cellular enzyme complex (METTL3/METTL14) which deposits m^6^A modifications. The METTL3/METTL14 complex is normally localized in the nucleus and deposits m^6^A modification co-transcriptionally (Huang et al. 2019). We determined whether METTL3/METTL14 function could be compromised during infection due to altered cellular localization. We observed that during SARS-CoV-2 infection METTL3 was partially relocalized from the nucleus to the cytoplasm (**Fig. 2F**). Importantly, cytoplasmic localization of METTL3 was more evident with the B.1 and B.1.1.7 variants compared to B.1.351 (**Fig. 2F-H**), which is consistent with a greater reduction in m^6^A level in B.1 and B.1.1.7 infections. To verify that the METTL3 antibody used was specific and did not detect a viral antigen in infected cells, we infected both control (Control-sh) and METTL3 knock-down (KD) Vero cells with the B.1 variant (**Supplemental Fig. S2G, S2H**). We observed a drastic reduction in METTL3 signal upon METTL3 KD in non-infected cells. After infection with B.1, the Control-sh cells showed METTL3 localization in both nuclear and cytoplasmic compartments, however, in the METTL3 KD condition, this staining was absent suggesting that the METTL3 antibody was indeed specific (**Supplemental Fig. S2I**). The nuclear localization of METTL14 was mostly unaffected during SARS-CoV-2 infection (**Supplemental Fig. S2J**). As m^6^A loss was marked in infected cells, we wondered if partial METTL3 cytoplasmic localization was sufficient to compromise METTL3/METTL14 functional complex formation. To test this, we performed a proximity ligation assay (PLA) to detect the METTL3/METTL14 complex in the non-infected and SARS-CoV-2 infected cells. We observed that the METTL3/METTL14 PLA signal was decreased in infected cells, with an effect that was more marked in B.1 and B.1.1.7 compared to B.1.351, where METTL3 localization was also less affected (**Fig. 2I, J**).

### SARS-CoV-2 genomic RNA contains m^6^A modification

Recently several studies reported the presence of m^6^A modification on SARS-CoV-2 genomic RNA (Li et al. 2021; Liu et al. 2021; Zhang et al. 2021b; Burgess et al. 2021). As we observed that the key m^6^A depositing enzyme METTL3 showed partial relocalization to the cytoplasm where SARS-CoV-2 genome replication occurs, we investigated the m^6^A profile in viral RNAs. Our strand-specific m^6^A RIP-seq data allowed us to profile m^6^A in both the positive (genomic) and the negative (replicative intermediates) strands of viral RNA for the three SARS-CoV-2 variants. Consistent with previous reports, we detected m^6^A enriched regions at several positions of the positive-strand SARS-CoV-2 genome for all three variants, with broader m^6^A peaks detected in the *N* gene region (Li et al. 2021; Liu et al. 2021; Zhang et al. 2021b; Burgess et al. 2021) (**Fig. 3A; Supplemental Data S3**). Using liquid chromatography-tandem mass spectrometry (LC-MS/MS), we then determined the presence of a relative number of m^6^A residues in the SARS-CoV-2 RNA genome. In these experiments, viral genomic RNA was isolated from infected Vero cell supernatant, and the ratio of A/m^6^A was measured using a synthetic m^6^A containing RNA oligo as a standard. Using the parallel reaction-monitoring (PRM) mode, we monitored LC-MS/MS profiles for m/z 268.0–136.0 and m/z 282.0–150.1, which corresponds to A and m^6^A respectively (Sun et al. 2021). We observed that each SARS-CoV-2 genome contains on average ten m^6^A modifications (**Supplemental Fig. S3A**), consistent with a recent report (Li et al. 2021). We observed m^6^A containing peaks were also present in the negative strand of the viral RNA, mainly in the N/ORF10 region (**Fig. 3B**). We then validated the presence of m^6^A in both positive and negative strands by strand-specific m^6^A RIP-qPCR for the B.1.1.7 variant (**Fig. 3C**). Further, using SARS-CoV-2 infected patient RNA samples from throat/nose swabs, we could detect m^6^A enrichment in the *N* gene region in both the positive and negative strands of SARS-CoV-2 RNAs (**Fig. 3D**). We verified upregulation of selected interferon-stimulated genes in SARS-CoV-2 infected patient throat/nose swab RNA as previously reported **(Supplemental Fig. S3B)** (Lorè et al. 2021; Gao et al. 2021). The presence of m^6^A peaks in both strands of SARS-CoV-2 RNA suggests an important functional role for such modifications (Liu et al. 2021; Li et al. 2021; Zhang et al. 2021b; Burgess et al. 2021). Inspection of publicly available viral-RNA protein interaction data shows that m^6^A reader protein partners are enriched in viral RNA interacting proteins (Schmidt et al. 2021). The m^6^A readers previously shown to interact with SARS-CoV-2 RNA were searched for known interacting protein partners in the STRING database. We found that these interacting proteins are enriched in RNA metabolism and viral process pathways (**Supplemental Fig. S3C, top panel**) and that these proteins show overlap with the SARS-CoV-2 RNA-protein interactome (**Supplemental Fig. S3C, bottom panel**). These observations suggest that SARS-CoV-2 may utilize m^6^A residues in its genome to recruit other RNA-binding proteins to the viral RNA using m^6^A readers as intermediates. This is consistent with the recently reported role of m^6^A reader YTHDF proteins in SARS-CoV-2 infection (Burgess et al. 2021). Given m^6^A has been implicated in other RNA viruses, detailed characterization of interactions between m^6^A readers and SARS-CoV-2 RNA will further elucidate the role of m^6^A reader proteins in host-virus interaction (Kim and Siddiqui 2021; Lichinchi et al. 2016).

**Fig. 3.**
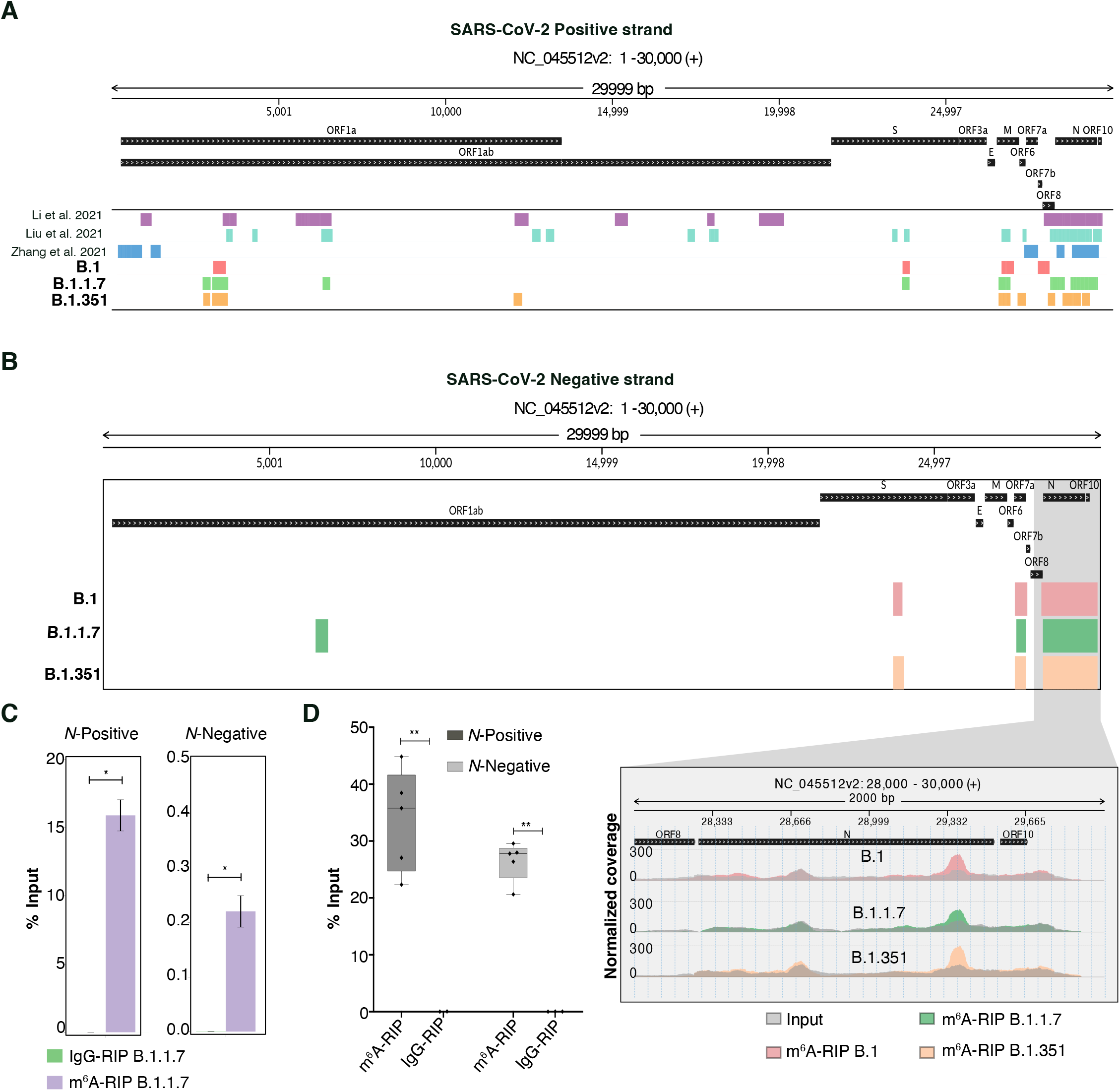
SARS-CoV-2 genomic RNA contains m^6^A modification. (A) Identified m^6^A peaks on SARS-CoV-2 positive strand from B.1, B.1.1.7 and B.1.351 variants are shown. The m^6^A peaks in the positive strand of SARS-CoV-2 RNA from publicly available data. (B) m^6^A peaks on the SARS-CoV-2 negative strand. The presence of m^6^A peaks in the *N* gene region is highlighted and normalized coverage is shown on the panel below. (C, D) m^6^A-RIP qPCR data showing enrichment of *N* gene region in both positive strand and the negative strand of viral RNA (C) 24 h post-infection with B.1.1.7 in Vero cells, data represented as a percentage of input, and IgG was used as negative control. Statistics: two-tailed paired *t*-test, n=3, *: p < 0.05 (D) in the infected patient samples (n=5). Data is represented as a percentage of input. IgG was used as a negative control. Statistics: two-tailed paired *t*-test, **: p < 0.01.

### m^6^A loss modulates cellular gene expression in SARS-CoV-2 infected cells

To better understand the effects of m^6^A modulation on cellular gene expression during viral infection, we first identified the set of transcripts showing a change in m^6^A level during infection with the three SARS-CoV-2 variants compared to non-infected cells. We then compared the genes with lost m^6^A (shows loss of at least one m^6^A peak compared to non-infected cells) or with gained m^6^A (no m^6^A peak present in non-infected cells, but one or more m^6^A peak gained after infection) with our set of differentially expressed genes (up- and down-regulated during infection). We found that the list of up- and down-regulated genes overlapped with that of genes with lost and gained m^6^A. In general, down-regulated genes showed higher overlap with lost m^6^A genes across the three variants, whereas the up-regulated genes were over-represented in the gained m^6^A gene category (**Fig. 4A**). To better understand how loss and gain of m^6^A contributed to gene expression, we performed a cumulative distribution function (CDF) analysis that measured RNA abundance using RNA-seq data in genes with either lost or gained m^6^A peaks. We found lost m^6^A genes showed a decrease in abundance whereas gained m^6^A genes were increased in abundance post-SARS-CoV-2 infection for the three variants studied (**Supplemental Fig. S4A**). Similar findings were obtained when genes with m^6^A gain, or loss were compared to genes with no change in m^6^A level (retained genes) (**Supplemental Fig. S4B**). We further categorized genes in non-infected cells based on m^6^A content, to either non-m^6^A, low (one peak), medium (two peaks), or high (three or more peaks) m^6^A. We observed higher m^6^A-containing genes were more susceptible to viral infection, with a decreased expression of these genes upon infection with SARS-CoV-2 variants (**Fig. 4B**). This suggests that m^6^A modification on these genes contributes to their expression and their expression is compromised due to the global loss of m^6^A during viral infection. We chose a few genes with varying levels of m^6^A in non-infected cells and validated their down-regulation during viral infection (**Fig. 4C)**. Expression of these genes was also decreased in METTL3 KD cells, indicating an m^6^A-dependent expression (**Supplemental Fig. S4C**). The exception was EID1, which did not change expression in absence of METTL3, consistent with this gene having no m^6^A modification (**Supplemental Fig. S4C**). Infection with the B.1.617.2 (Delta) variant also down-regulated the expression of the five tested genes (**Fig. 4C**). We checked the localization of METTL3 after B1.617.2 infection and observed that METTL3 was partially relocalized from the nucleus to the cytoplasm in B.1.617.2 infection as well (**Supplemental Fig. S4D**). We then performed a pathway analysis for the up- and down-regulated genes with lost m^6^A. We observed that down-regulated genes with lost m^6^A were mostly involved in cilium organization, whereas up-regulated genes with lost m^6^A were primarily part of pathways related to covalent chromatin/histone modifications (**Supplemental Fig. S4E**). We validated the down-regulation of some of the known cilia-related genes such as *HDAC6* and *PKHD1* (Ran et al. 2015; Zhang et al. 2004) **(Fig. 4C**) consistent with a recent report suggesting widespread loss of motile cilia after SARS-CoV-2 infection (Robinot et al. 2021). The genes with lost m^6^A involved in covalent chromatin modification and up-regulated during infection, such as KDM6A (**Supplemental Data S2**), were recently identified as proviral genes in a CRISPR screen (Wei et al. 2021b). Thus, our data suggest that loss of m^6^A during SARS-CoV-2 infection establishes a pattern of host cell gene expression that favors viral replication.

**Fig. 4.**
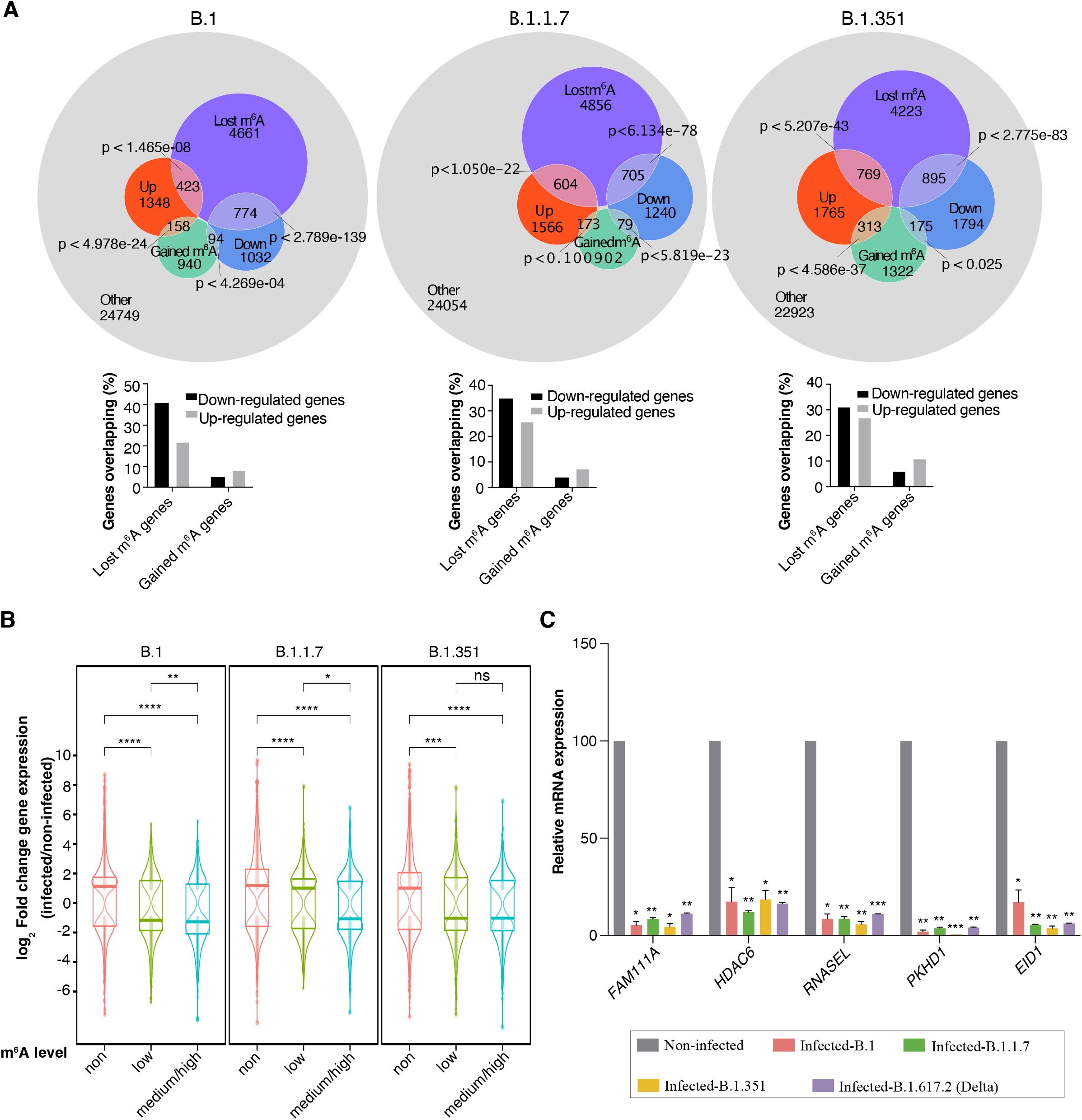
Gene expression post-viral infection and global m^6^A loss. (A, upper panel) Overlap diagrams of differentially expressed and m^6^A - modified genes after infection. Each set shows the number of up-regulated and down-regulated genes associated with lost or gained m^6^A modifications. The remaining genes are shown in gray. Hypergeometric test p-values depict the calculated probability of overlap between differentially expressed and m^6^A lost/gained genes. (A, lower panel) Bar-graph summarizing the percentage of up-/down-regulated genes overlapping with either lost or gained m^6^A genes. (B) Log_2_ fold-change distributions of differentially expressed genes post-infection with three different variants, categorized according to their m^6^A level: non, low, medium/high. Statistical significance was calculated using the Wilcoxon test. ns: non-significant, *: p < 0.05, **: p < 0.01, ***: p < 0.001, ****: p < 0.0001. (C) Relative mRNA expression of genes with varying levels of m^6^A (*FAM111A. PKHD1-* high m^6^A; *HDAC6*-medium m^6^A; *RNASEL-low* m^6^A; *EID1*-no m^6^A) post-viral infection with variant indicated compared to non-infected cells. qPCR data was normalized to *TBP* and *POL2RG*. Statistics: two-tailed *t*-test; *: p < 0.05, **: p < 0.01, ***: p < 0.001, n=2.

m^6^A modification is known to be implicated in alternative splicing (Wei et al. 2021a). Considering the global loss of m^6^A during SARS-CoV-2 infection, we looked for differential exon use in the RNA-seq data. We identified about 790 differential exon usage (DEU) events common to cells infected with the three SARS-CoV-2 variants (**Supplemental Data S4**). For instance, we observed that exon 1 of the *COL6A2* collagen gene is differentially included during infection, which is associated with the loss of m^6^A peaks surrounding the newly included exon1 (**Supplemental Fig. S4F**). We also found that DEU-containing genes had on average more m^6^A peaks compared to random m6A-positive genes in non-infected cells (**Supplemental Fig. S4G**). Further, we observed that m^6^A containing DEU genes showed a loss of m^6^A peaks during infection (**Supplemental Fig. S4H**).

### Treatment with Selinexor restores METTL3 cellular localization during SARS-CoV-2 infection

During the SARS-CoV-2 infection, we observed partial relocalization of METTL3 to the cytoplasm from its original nuclear location. Considering that m^6^A modification of cellular mRNA is deposited co-transcriptionally in the nucleus (Huang et al. 2019), we aimed to restore METTL3 nuclear localization to see if we could antagonize the effects of SARS-CoV-2 infection. We observed that the expression of *XPO1* (or Exportin 1), one of the major nuclear export proteins, was up-regulated during infection (**Fig. 5A; Supplemental Fig. S5A**). Therefore, we set to determine whether XPO1 was involved in the relocalization of METTL3 during viral infection. Using PLA, we observed that XPO1 and METTL3 interacted with each other in Vero cells (**Fig. 5B**). We then treated these cells during infection with Selinexor, a well-characterized XPO1 inhibitor (Kashyap et al. 2021). We observed that Selinexor treatment resulted in the restoration of METTL3 localization to the nucleus of B.1 infected cells (**Fig. 5C, D**). We then checked if the restoration of METTL3 localization could rescue METTL3/METTL14 complex formation, which was compromised in infected cells. We observed that the METTL3/METTL14 PLA signal was robustly detected in B.1 and B.1.1.7 infected cells treated with Selinexor, but not in infected cells treated with DMSO (**Fig. 5E; Supplemental Fig. S5B**). These results suggested that change in METTL3 localization perturbed the formation of the METTL3/METTL14 complex and that promoting METTL3 nuclear localization by Selinexor was sufficient to drive the formation of the METTL3/METTL14 complex in infected cells. Importantly, Selinexor treatment was also effective in decreasing SARS-CoV-2 infection, as measured by Spike protein positivity for all the four variants tested (B.1, B.1.1.7, B.1.351, B.1.617.2), without decreasing cell viability (**Supplemental Fig. S5C, D**). Of note, the efficacy of Selinexor in preventing SARS-CoV-2 infection has also been reported recently (Kashyap et al. 2021).

**Fig. 5.**
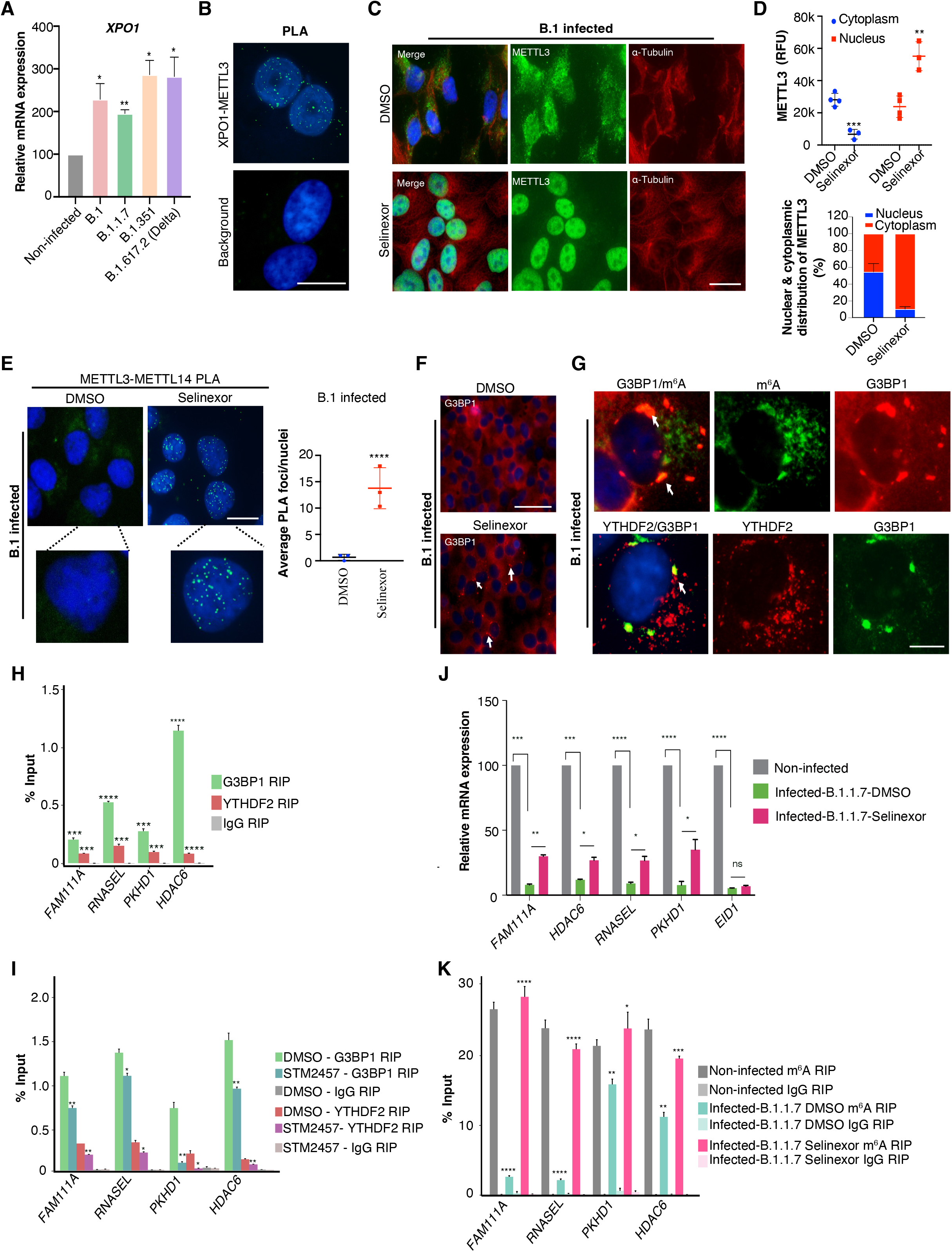
Treatment with Selinexor restores METTL3 cellular localization during viral infection. (A) Relative expression of the *XPO1* gene evaluated post-SARS-CoV-2 infection (variants as specified) using RT-qPCR. Data was normalized to *TBP* and *POL2RG*. Two-tailed *t*-test; *: p < 0.05, **: p < 0.01, n=3. (B) Top panel: Proximity ligation assay (PLA) in non-infected Vero cells depicting interacting XPO1 and METTL3 foci, (bottom panel) background control for PLA with only XPO1 antibody. Scale bar is 20 μm. (C-D) Rescue of METTL3 localization: (C) METTL3 and α-Tubulin co-immunostaining in Vero cells post-infection (24 h) with B.1 SARS-CoV-2 that were either treated with DMSO or Selinexor (150nM). Scale bar is 20 μm. (D, top panel) METTL3 intensities (RFU) from the immunostaining performed in (C). METTL3 (RFU) in the nucleus and cytoplasm were estimated using ImageJ with help of DAPI marking the nucleus and α-Tubulin staining the cytoplasm. Data are shown as Mean ± SD. Data presented from multiple experiments with number of cells counted, n=<70. One-way ANOVA test was performed, **: p < 0.01, ***: p < 0.001. (D, bottom panel) Percentage of METTL3 distribution in nucleus and cytoplasm calculated from intensities from the upper panel. (E, top panel) PLA depicts interacting METTL3 and METTL14 foci in the nucleus (marked by DAPI) of Vero cells infected with B.1 SARS-CoV-2 variants and treated with either DMSO or Selinexor (150nM). Scale bar is 20 μm. (Bottom panel). (E, bottom panel) Quantification of METTL3-METTL14 PLA foci as detected in the above panel. The number of PLA foci/nuclei are shown as Mean ± SD. Data presented from multiple experiments with number of cells counted, n=>100. One-way ANOVA test was performed, ****: p < 0.0001. (F) Immunostaining showing G3BP1 localization after DMSO or Selinexor (150 nM) treatment in B.1-infected cells (24-h post-infection). White arrow highlights some of the G3BP1 foci. Scale bar is 50 μm. (G) Immunostaining showing G3BP1 and m^6^A/YTHDF2 localization after Selinexor treatment in B.1-infected cells (24-h post-infection). White arrow highlights some of the G3BP1 foci overlapping with either m^6^A or YTHDF2. Scale bar is 10 μm. (H) G3BP1 and YTHDF2 RIP showing enrichment of m^6^A positive genes in Vero cells. IgG was used as negative control. Statistics: two-tailed *t*-test; ***: p < 0.001, ****: p < 0.0001. n=4. (I) G3BP1 and YTHDF2 RIP showing enrichment of m^6^A positive genes in Vero cells treated with either METTL3 inhibitor- STM2457 or DMSO. IgG was used as negative control (two-tailed *t*-test; *: p < 0.05, **: p < 0.01, n=4). (J) Relative mRNA expression of genes with varying levels of m^6^A after Selinexor treatment in B.1.1.7 infected cells. Treatment with Selinexor partially restores the expression of m^6^A-modified genes (*FAM111A, HDAC6, RNASEL*, and *PKHD1*), but not the non-m^6^A gene (*EID1*). qPCR data was normalized to *TBP* and *POL2RG*. Statistics: two-tailed *t*-test; *: p < 0.05, **: p < 0.01, ***: p < 0.001, ****: p < 0.0001), n=3. (K) m^6^A-RIP qPCR demonstrating recovery of m^6^A levels at selected genes in Selinexor treated B.1.1.7 infected cells. DMSO was used as treatment control and IgG was used as negative control for RIP. Statistics: two-tailed *t*-test; *: p < 0.05, **: p < 0.01, ***: p < 0.001, ****: p < 0.0001 n=3).

m^6^A-modified RNA and m^6^A reader YTHDF proteins have been implicated in stress granule formation by promoting the recruitment of stress granule proteins such as G3BP1 (Fu and Zhuang 2020). Stress granule formation is known to be compromised in SARS-CoV-2 infected cells, and knock-down of the stress granule protein G3BP1 enhances SARS-CoV-2 infection (Zheng et al. 2021). We reasoned that the inactivation of stress granule formation by SARS-CoV-2 could be mediated by the global loss of m^6^A in cellular mRNAs that we uncovered in infected cells. Therefore, we tested if Selinexor-mediated reversal of METTL3 nuclear localization could promote stress granule formation in infected cells by reinstalling m^6^A modification in cellular mRNAs. We observed the appearance of stress granules in Selinexor treated, but not in DMSO-treated (control), SARS-CoV-2 infected cells (**Fig. 5F; Supplemental Fig. S5E**). Immunostaining showed that stress granules in Selinexor-treated infected cells often overlapped with m^6^A-modified RNAs and with the m^6^A reader protein YTHDF2 (**Fig. 5G; Supplemental Fig. S5F**). We then used an RNA immunoprecipitation (RIP) assay to check the interaction of four validated mRNAs which show loss of m^6^A and down-regulation post-infection with the m^6^A reader YTHDF2 and the stress granule protein G3BP1. Indeed, we found that these mRNAs coprecipitated with both G3BP1 and YTHDF2 (**Fig. 5H**). Inhibiting the METTL3 enzyme by the small molecule inhibitor STM2457 (Yankova et al. 2021) decreased the interaction of the four mRNAs with G3BP1 and YTHDF2, suggesting that m^6^A is required for such interaction (**Fig. 5I**). Treatment with Selinexor, which restored METTL3 nuclear localization during infection, also rescued the expression of the four candidate down-regulated genes, which included the HDAC6 and PKHD1 genes related to cilia formation (**Fig. 5J**). We also observed that Selinexor treatment could reinstate the m^6^A modifications that were lost during infection in these four mRNAs (**Fig. 5K**). Therefore, rescuing nuclear localization of METTL3 by Selinexor promoted stress granule formation in infected cells, restored cellular gene expression, and inhibited SARS-CoV-2 infection *in vitro*, highlighting the possibility of targeting the m^6^A pathway as an antiviral strategy.

### Global loss of m^6^A modification in primary human airway epithelial cells upon SARS-CoV-2 infection

Next, we aimed to validate the global loss of m^6^A methylation during SARS-CoV-2 infection in human cell infection models. To this goal, we tested METTL3 localization post-SARS-CoV-2 infection in bronchial epithelial cell line, BEAS-2B. BEAS-2B cells were positive for spike protein and METTL3 showed partial relocalization from the nucleus to the cytoplasm in these infected cells (**Fig. 6A; Supplemental Fig. S6A**). Further, we also tested METTL3 localization post-infection in primary human bronchial epithelial cells grown in monolayer culture and similar to BEAS-2B we observed partial cytoplasmic relocalization of METTL3 (**Supplemental Fig. S6B, S6C**).

**Fig. 6:**
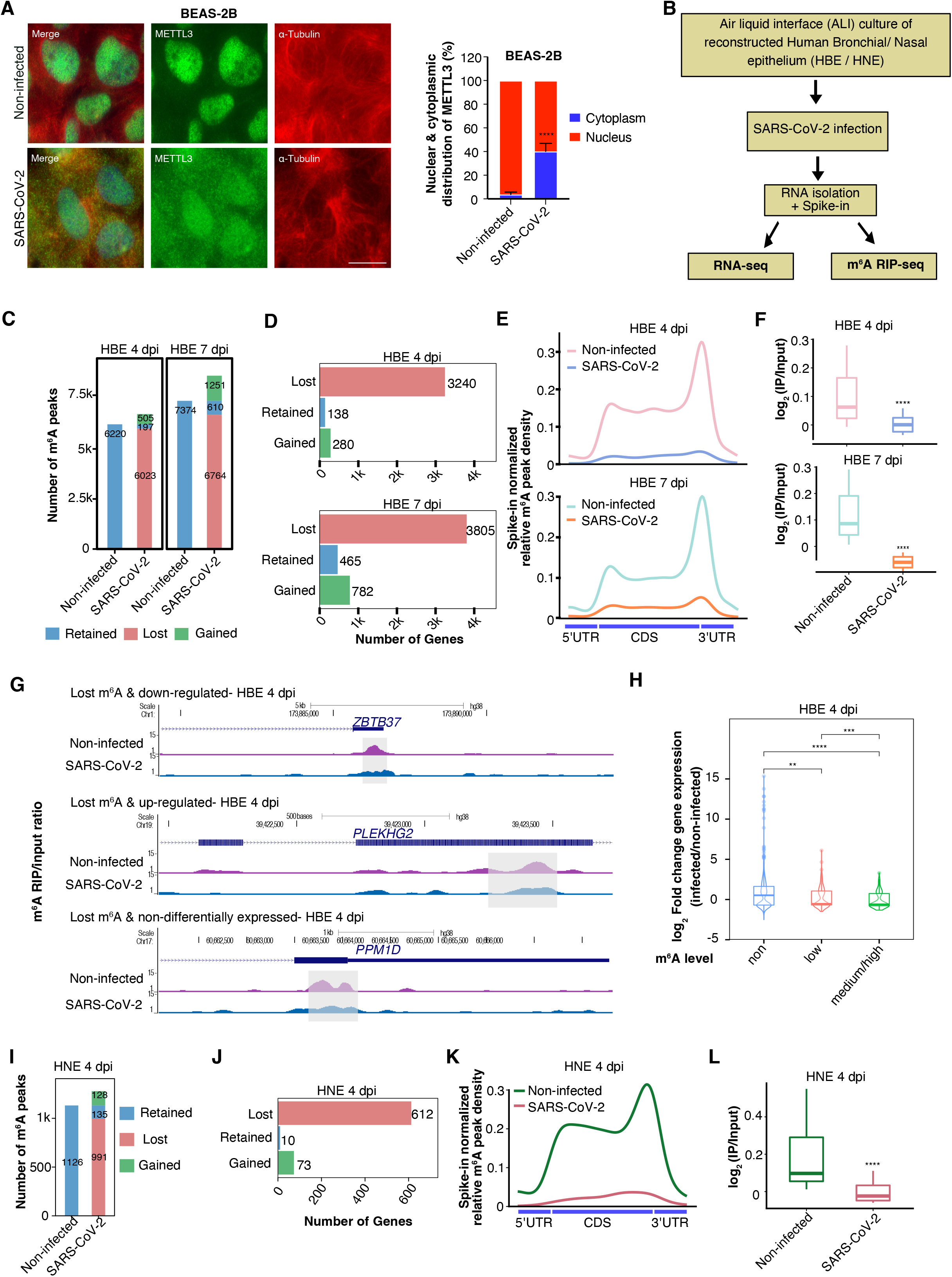
Loss of m^6^A in human airway epithelia after SARS-CoV-2 infection. (A, left panel) METTL3 localization in BEAS-2B cells post-infection with SARS-CoV-2. The scale bar is 20 μm. (A, right panel) The percentage distribution of METTL3 in the nucleus and cytoplasm was calculated using ImageJ with help of DAPI marking the nucleus and α-Tubulin staining the cytoplasm. Data are shown as Mean ± SD. Data presented from multiple experiments with the number of cells counted, n=< 300. Unpaired *t*-test; ****: p < 0.0001 (B) Flow chart describing the experimental design for SARS-CoV-2 infection of reconstructed Human Bronchial /Nasal Epithelium at ALI. (C) Total number of m^6^A peaks classified as retained, gained, or lost, after 4- or 7- days post infection (dpi) with SARS-CoV-2, compared to non-infected HBE. (D) Bar plots representing the number of genes that lost, gained, or retained m^6^A in HBE after 4- and 7- dpi. (E) Metagene plots showing spike-in normalized relative m^6^A peak density distribution at host genes in non-infected and SARS-CoV-2 infected HBE cells 4- and 7-dpi. (F) Boxplot comparing log_2_ ratio of m^6^A signal over input at lost m^6^A peak regions (± 125bp from m^6^A peak submit) obtained from panel (C) at 4- and 7- dpi in HBE. Statistical significance was calculated using the Wilcoxon test. ****: p < 0.0001. (G) Genome browser screenshots demonstrating spike-in normalized m^6^A (IP/input ratio) tracks for three different genes in non-infected and SARS-CoV-2 infected HBE 4- dpi. m^6^A peak region identified using MACS peak caller in non-infected cells are highlighted using the gray box. (H) log_2_ fold-change distributions of differentially expressed genes in HBE 4- dpi, categorized according to their m^6^A level: non, low, medium/high. Statistical significance was calculated using the Wilcoxon test. **: p < 0.01, ***: p < 0.001, ****: p < 0.0001. (I) Total number of m^6^A peaks classified as retained, gained, or lost, after 4- dpi with SARS-CoV-2, compared to non-infected HNE. (J) The number of genes that lost, gained, or retained m^6^A in HNE post-SARS-CoV-2 infection. (K) Metagene plots showing spike-in normalized relative m^6^A peak density distribution in non-infected and SARS-CoV-2 infected HNE. (L) Boxplot comparing log_2_ ratio of m^6^A signal over input at lost m^6^A peak regions (± 125bp from m^6^A peak submit) obtained from panel (I) at 4 dpi in HNE. Statistical significance was calculated using the Wilcoxon test. ****: p < 0.0001.

We then used a reconstructed human bronchial epithelium (HBE) as the SARS-CoV-2 infection model, where cells are differentiated by the culture at the air/liquid interface (ALI) forming a pseudostratified ciliated epithelium prior to viral infection as described previously (Robinot et al. 2021). RNA was extracted from HBE at 4- and 7- days post-infection (dpi) and we observed high levels of SARS-CoV-2 replication as measured by *N* gene expression (**Supplemental Fig. S6D**). Extracted RNA from HBE at 4- and 7- dpi was supplemented with spike-in bacterial RNA and used in m^6^A RIP-seq experiments (**Fig. 6B**). SARS-CoV-2 reads contributed 0.48 and 0.47% of the total reads at 4- and 7- dpi respectively (**Supplemental Fig. S6E**). Consistent with our observations in infected Vero cells, we observed a marked loss in the number of m^6^A peaks post-infection (**Fig. 6C; Supplemental Fig. S6F; Supplemental Data S3**). The top enriched motif in the m^6^A peaks from uninfected HBE contained the “GGAC” sequence, whereas the top motif was more degenerate in infected HBE (**Supplemental Fig. S6G**). A few transcripts showed a gain of m^6^A during infection in the HBE model, but the fraction of transcripts with gained m^6^A peaks was much lower compared to that in Vero cells (**Fig. 6C)**. We observed a drastic decrease in the number of m^6^A-positive genes in HBE on both 4- and 7- dpi (**Fig. 6D, E)**. m^6^A enrichment (IP/input) and spike-in normalized m^6^A signal over the lost peaks were significantly reduced in the SARS-CoV-2 infected cells, while input RNA was unchanged over these lost peaks (**Fig 6F; Supplemental Fig. 6H**). Genome browser visualization of three chosen genes showed the loss of m^6^A peaks at 4- dpi (**Fig. 6G**). We could detect m^6^A peaks in both the positive and negative strands of viral RNA at 4- dpi. We also detected m^6^A peaks in the positive strand of viral RNA at 7- dpi, but m^6^A peaks were absent in the negative strand at day 7, possibly because the infection reaches a plateau or has already started to decrease at this stage, as described previously (Robinot et al. 2021) (**Supplemental Fig. S6I; Supplemental Data S3**). Genes related to the defense response to viruses and type I interferon signaling pathways were up-regulated upon SARS-CoV-2 infection in HBE, whereas genes related to cilium organization were the most down-regulated (**Supplemental Fig. S6J; Supplemental Data S1, S2**). In the HBE infection model, genes containing one or more m^6^A peaks were prone to down-regulation post-infection, similar to the findings obtained in Vero cells (**Fig. 6H**). Down-regulated genes with lost m^6^A were again enriched in the pathways of cilium organization and cilium assembly (**Supplemental Fig. S6K**), suggesting that m^6^A loss could be a major contributor to the loss of cilia as previously reported in the HBE model (Robinot et al. 2021). These data collectively confirm that SARS-CoV-2 infection causes a general perturbation of m^6^A-dependent gene expression in human primary epithelial cells.

We further evaluated changes in cellular m^6^A level following SARS-CoV-2 infection in reconstructed human nasal epithelia (HNE) cultures in ALI conditions, which also support robust SARS-CoV-2 infection (**Fig. 6B**) (Samelson et al. 2022). We verified SARS-CoV-2 infection in HNE 4- dpi using the expression of the *N* gene (**Supplemental Fig. S7A**). SARS-CoV-2 reads contributed 0.79% of total reads in the HNE (**Supplemental Fig. S7B**). Infected HNE showed a drastic loss of m^6^A peaks 4- dpi and a decrease in the number of m^6^A positive genes as well (**Fig. 6I-K**). As observed in HBE, m^6^A enrichment (IP/input) and spike-in normalized m^6^A signal over the lost peaks were significantly reduced in HNE, while input RNA was unchanged (**Fig 6F; Supplemental Fig. 6H**). (**Fig 6L; Supplemental Fig. S7C**). Importantly infected HNE showed reduced level of nuclear METTL3 with partial relocalization to the cytoplasm (**Supplemental Fig. S7D, S7E**). We observed that infected HNE were positive for spike protein and showed disturbed staining of cilia marker β-IV-Tubulin as reported earlier (Robinot et al. 2021) (**Supplemental Fig. S7D**). Taken together, our observations in multiple transformed and primary cell culture models suggest that SARS-CoV-2 infection causes a global loss of m^6^A in cellular RNA.

## DISCUSSION

We report that SARS-CoV-2 infection causes a global loss of m^6^A in cellular RNAs while enabling m^6^A addition in viral RNAs, highlighting how this virus usurps mRNA modification pathways to promote its replication. Mechanistically, we found that SARS-CoV-2 infection induced a partial relocalization of the m^6^A methyltransferase METTL3 from the nucleus to the cytoplasm, which compromised the formation of the METTL3/METTL14 complex in the nucleus and could account for the overall decrease in m^6^A modification in infected cells. The latter findings are consistent with those of Zhang et al., who reported a cytoplasmic localization of both METTL3 and METTL14 along with other m^6^A reader proteins in SARS-CoV-2 infected Vero cells (Zhang et al. 2021b). The extent of METTL3 cytoplasmic relocalization and its effect on METTL3/METTL14 complex function and m^6^A levels depended on the viral variants. The B.1 and B.1.1.7 variants induced a more pronounced METTL3 relocalization and a more profound loss of m^6^A than the B.1.351 variant, suggesting that differences in variant fitness may not only depend on their capacity to escape from the innate and adaptive immune responses (Alefishat et al. 2022), but also, on their capacity to perturb gene expression in infected cells. We also observed global loss of m^6^A peaks in primary human airway epithelial cells during SARS-CoV-2 infection, confirming the generality of our findings. In previous reports, however, a global loss in m^6^A was not detected in the Huh7 and A459 cell lines following SARS-CoV-2 infection, even though a decrease in m^6^A peaks was observed near the stop codon region in Huh7 infected cells (Liu et al. 2021; Burgess et al. 2021). The differential effect of SARS-CoV-2 infection on m^6^A regulation in Huh7 and A549 cells compared to Vero, HBE and HNE infection models may result from different infection rates, and/or different tissue origin of the cell types used (Saccon et al. 2021). Though we observed a global decrease in m^6^A peaks in infected cells, a fraction of transcripts nevertheless showed a gain in m^6^A peaks upon infection. These gained m^6^A modifications may be deposited by residual METTL3/METTL14 in the nucleus or by METTL16, which was recently described to also have m^6^A writing capacity in mRNAs (Su et al. 2022). Future investigations will be needed to understand how the variant-specific effect on m^6^A contributes to viral adaptation and how a subset of m^6^A peaks can be gained during infection.

Gene expression analysis also highlighted common pathways dysregulated by all the SARS-CoV-2 variants tested. We observed in particular that genes associated with RNA catabolism tended to be up-regulated in infected Vero cells. Several genes associated with the m^6^A pathway, including m^6^A readers and m^6^A erasers, were deregulated as well during infection. Change in the RNA splicing pattern of m^6^A-related genes was recently reported after acute depletion of METTL3, which was interpreted as a feedback loop to compensate for the acute depletion of m^6^A (Wei et al. 2021a). In SARS-CoV-2 infected cells, depletion of m^6^A following METTL3 relocalization and loss of functional METTL3/METTL14 complex in the nucleus probably activates a similar feedback loop, which could explain why genes involved in RNA catabolism and m^6^A modification were preferentially deregulated. Our combined analysis of differentially expressed genes and change in m^6^A peak abundance suggests that, in general, loss of m^6^A peaks correlates with decreased RNA expression. However, it is important to consider that loss of m^6^A was more widespread than mRNA decrease in the infected cells, and thus that loss of m^6^A did not always lead to decreased gene expression. Further investigation is needed to decipher the mechanisms that dictate how loss of m^6^A influences gene expression during viral infection. Though we have pointed out DEU events in the subset of genes with m^6^A loss, further study is required to understand the effect of m^6^A loss on the other cellular genes that show no change in expression. We observed that m^6^A containing genes were more prone to down-regulation during infection in both Vero and HBE infection models. The SARS-CoV-2 Nsp1 protein has been shown to inhibit nuclear export of cellular RNAs during infection and to promote host mRNA decay (Burke et al. 2021; Zhang et al. 2021a). The nuclear export of cellular RNA is known to be regulated by m^6^A in an YTHDC1- and Nuclear RNA Export Factor 1- (NFX1) dependent manner (Roundtree et al. 2017). Viral protein Nsp1 also inhibits NFX1 function, resulting in the retention of cellular mRNAs in the nucleus during SARS-CoV-2 infection (Zhang et al. 2021a). It will be interesting to determine if the loss of m^6^A and Nsp1-mediated inhibition of NFX1 act synergistically to cause nuclear retention of cellular mRNAs during infection, and if mRNAs retained in the nucleus might be more prone to rapid degradation.

Using a strand-specific m^6^A RIP-seq approach, we found that both the viral genomic RNA (positive strand) and replicative negative-strand RNAs carried m^6^A modification. In the human bronchial epithelium model, we detected a robust m^6^A peak signal in genomic RNA at 4- and 7- dpi. The m^6^A peaks were also present in the replicative negative strand at 4- dpi, but not at day 7, presumably because viral replication had already abated at this time point. In a previous report by Liu et al., m^6^A peaks could be detected on negative-strand of SARS-CoV-2 using m^6^A RIP-seq, but not with the m^6^A-CLIP (Cross-Linking and Immunoprecipitation) technique. The authors suggested lack of m^6^A detection could be due to the limited coverage of the negative strand in CLIP data (Liu et al. 2021). Though our data from Vero and human bronchial epithelium infection models suggest the presence of m^6^A peaks in the negative strand, future experiments using m^6^A-CLIP in the human cell infection model will be required to precisely identify the specific residues modified by m^6^A. Viral RNA m^6^A modification is known to have a pro-viral effect by allowing escape from RIG-I binding and limiting the induction of inflammatory gene expression (Li et al. 2021). As RIG-I is known to be activated by viral double-stranded replicative intermediates (Yamada et al. 2021) it will be interesting to check if the m^6^A modifications detected in the replicative negative strand of SARS-CoV-2 contribute to an escape mechanism from RIG-I. Though the majority of m^6^A peaks were similar across the three SARS-CoV-2 variants studied, there were m^6^A peaks that were specific to particular variants. Further detailed studies will be required to understand if the differences in m^6^A modification across variants contribute to changes in pathogenicity by differential recruitment of m^6^A reader proteins or by modulating the RIG-I-dependent escape mechanisms described above.

To explore new therapeutic options against SARS-CoV-2 infection, we have exploited the altered localization of the m^6^A writer METTL3. We succeeded in restoring METTL3 localization during infection by inhibiting the nuclear export protein XPO1 with Selinexor. Selinexor treatment not only restored METTL3 localization but also increased the expression of specific genes which showed down-regulation with concomitant loss of m^6^A during infection. XPO1 inhibition by Selinexor was previously proposed to be effective in limiting SARS-CoV-2 infection by promoting the nuclear relocalization of the ACE-2 receptor (Kashyap et al. 2021). Our study provides a further mechanistic explanation for the effectiveness of XPO1 inhibition during SARS-CoV-2 infection. Though we provide evidence that altered METTL3 localization could be mediated by XPO1, we do not rule out other possible mechanisms which might synergistically affect METTL3 localization during SARS-CoV-2 infection. In particular, cytokines that are secreted by infected cells might influence METTL3 localization in both infected and bystander cells, resulting in broad perturbations in cellular m^6^A, a notion that requires further investigation. Cellular m^6^A-modified RNAs and m^6^A reader YTHDF proteins promote stress granule condensate formation, which contributes to the antiviral cellular defense (Fu and Zhuang 2020). The SARS-CoV-2 N-protein has been shown to phase-separate with the stress granule protein G3BP1 and thereby prevent stress granule formation in infected cells (Wang et al. 2021). We propose an additional mechanism for the inhibition of stress granule formation through the depletion of m^6^A on cellular RNA. The recovery of stress granules formation in Selinexor-treated infected cells may indeed result from restored m^6^A levels in cellular RNA, which could be visualized by the co-localization of G3BP1, m^6^A, and YTHDF2. We observed that Selinexor treatment also had functional consequences, by restoring the gene expression and m^6^A enrichment during infection. The recovery of stress granules may have also contributed to stabilizing these RNAs, as association with stress granule proteins such as G3BP1 has been shown to regulate the stability of cellular RNAs (Laver et al. 2020; Somasekharan et al. 2020). Some of the validated m^6^A-dependent RNAs showed an association with G3BP1 and YTHDF2, suggesting that these RNAs could indeed be guided to stress granules in an m^6^A-dependent manner.

Collectively, our findings highlight how SARS-CoV-2 perturbs the m^6^A RNA modification pathway to deregulate cellular RNAs and limit stress granule formation. Change in the cellular localization of METTL3 in infected cells resulted in a global loss of m^6^A in cellular mRNAs, whereas viral RNAs remained m^6^A modified. We propose that rescuing METTL3 localization during infection could be explored as a novel antiviral strategy against SARS-CoV-2.

## METHODS

### Cell culture, SARS-CoV-2 infection and XPO1 inhibitor treatment

SARS-CoV-2 was isolated from samples with viral genomes sequenced at the diagnostic laboratory (Ringlander et al. 2021). Isolates from the B.1, B.1.1.7, B.1.351, and B.1.617.2 pango lineages were used to infect Vero CCL-81 cells (ATCC) grown in DMEM, 1% penicillin-streptomycin and 2% fetal calf serum at 37°C and in 5% CO_2_. B.1 is the lineage with the D614G spike amino acid change. B.1.1.7 is the Alpha variant (WHO label) and the earliest documented variant in the UK (Sep-2020). B.1.1.351 is the Beta variant (WHO label) and was first documented in South Africa (May 2020). T25 flasks with Vero CCL-81 (ATCC) were infected with the three different SARS-CoV-2 variants (1000 TCID50) and medium or infected cells were harvested in TRIzol 48 h post-infection for RNA-sequencing studies. In all cases, the amount of infection was verified by immunostaining and RT-qPCR (see method below). Vero CCL-81 cells were cultured in chamber slides and the different SARS-CoV-2 variants were diluted in medium to 1000 TCID_50_ (DMEM with 1% penicillin-streptomycin and 2% fetal calf serum), supplemented with DMSO or Selinexor (150nM). 200 μl of the diluted virus was added to each well and the slides were incubated for 24 h at 37°C at 5% CO_2_. The cells were fixed using 4% formaldehyde in PBS for 10 min at room temperature (RT) in chamber slides and washed three times in PBS.

Primary human bronchial epithelial cells (Lonza, CC-2540S) and bronchial epithelial cell line BEAS-2B (ATCC) were received as a gift from Professor Madeleine Rådinger group, Gothenburg University, Sweden, and were cultured in monolayer following manufacturer instructions. Primary human bronchial epithelial and BEAS-2B cells were infected with the B.1.1.7 variant of SARS-CoV-2 in a similar way to Vero cells as described above in their corresponding culture media and fixed using 4% formaldehyde in PBS for 10 min at RT in chamber slides 72 h post-infection.

### SARS-CoV-2 infection of reconstructed human bronchial and nasal epithelia

MucilAir™ cultures, corresponding to reconstructed human bronchial and nasal epithelia (HBE and HNE, respectively) differentiated at the air/liquid interface (ALI), were purchased from Epithelix (Saint-Julien-en-Genevois, France). Cultures were maintained in ALI conditions in transwells with 700 μL of MucilAir™ medium (Epithelix) in the basal compartment and kept at 37°C under a 5% CO_2_ atmosphere. SARS-CoV-2 infection was performed in epithelial cultures as previously described (Robinot et al. 2021). Briefly, the apical side of ALI cultures was washed once, and cells were then incubated with the isolate BetaCoV/France/IDF00372/2020 (EVAg collection, Ref-SKU: 014V-03890) diluted in 150 μl of DMEM medium. The viral input was left on the apical side for 4 h at 37 °C, and then removed by 3 apical washes. Positivity for SARS-CoV-2 infection was tested by RT-qPCR as described below.

### Immunofluorescence Staining

Cells were fixed with 4% formaldehyde for 10 min, followed by two washes with phosphate-buffered saline (PBS). Then cells were permeabilized using 0.25% Triton X-100 in PBS for 10 min followed by two washes with PBS-0.1% Tween 20 (PBST). Blocking was performed in 3% bovine serum albumin (BSA) in PBST for an hour at RT. The cells were incubated overnight at 4 °C with the following primary antibodies: anti-METTL3 rabbit antibody (1:300, ab195352; Abcam), anti-METTL14 rabbit antibody (1:300, HPA038002; Atlas Antibodies), anti-α-Tubulin mouse antibody (1:1000, T5168; Merck), anti-SARS-CoV-2 Spike glycoprotein rabbit antibody (1:300, ab272504; Abcam), anti-G3BP mouse antibody (1:500, ab56574; Abcam), anti-m^6^A rabbit antibody (1:400, 202003, Synaptic Systems), and anti-YTHDF2 rabbit antibody (1:100, ab246514; Abcam). The slides were washed three times for 5 min in PBST and subsequently incubated with secondary antibodies conjugated with Alexa Fluor 488 and Alexa Fluor 555 fluorochromes (1:800; Invitrogen), for an hour in the dark at RT. After incubation with the secondary antibodies, cells were washed three times for 5 min in PBST. Prolong Gold with DAPI (Thermo Fisher Scientific–Life Technologies) was added to each coverslip and air-dried in the dark to detect nuclei. Slides were imaged in a fluorescence microscope (EVOS™ FL Auto, Thermo Fisher Scientific) using 20X and 60X oil immersion objectives by keeping the same parameter during image acquisition.

For immunofluorescence staining in HNE, MucilAir™ cultures were fixed in 4% PFA for 30 mins at RT, washed two times with PBS, and cut using a scalpel blade. Membrane pieces were placed in 10 μl drops on parafilm, and subsequent staining steps were performed on parafilm at RT. Cells were permeabilized with 0.5% Triton X-100 in PBS for 20 min and then blocked in 0.1% Tween 20, 1% bovine serum albumin (BSA), 10% fetal bovine serum, 0.3 M glycine in PBS for 30 min. Samples were incubated overnight at 4°C with AF647 conjugated rabbit anti-METTL3 primary antibody (ab217109; Abcam; 1:100 dilution), AF488 conjugated rabbit anti-β-IV-Tubulin (ab204003; Abcam; 1:250 dilution) and anti-spike (1:100, is a gift from Hugo Mouquet (Pasteur Institute). The samples were washed thrice in PBS followed by one hour incubated at RT with secondary antibody AF555-conjugated goat anti-mouse (1:500 dilution). Samples were counterstained with Hoechst and mounted in FluoromountG (Thermo Fisher Scientific) before observation with a Leica TCS SP8 confocal microscope (Leica Microsystems).

For quantification of METTL3 localization, images were analyzed with ImageJ. Fluorescence channels corresponding to the nuclear marker (DAPI/ Hoechst) and cytoplasmic marker (anti-α-Tubulin or anti-β-IV-Tubulin) were split to mark the nuclear and cytoplasmic border followed by quantification of METTL3 intensity (Relative Fluorescence Unit: RFU) in nucleus and cytoplasm.

### RNA-seq and m^6^A RNA Immunoprecipitation sequencing (m^6^A RIP-seq)

RNA was isolated from both SARS-CoV-2 infected (48h post-infection) and non-infected (mock) Vero cells using TRIzol reagent (Thermo Fisher Scientific, 15596026) and Direct-zol RNA Miniprep (ZYMO research, R2050). For RNA extraction of primary epithelial cultures, cells were washed in cold PBS and then lyzed in 150 μl of TRIzol reagent (Thermo Fisher Scientific) added to the apical side of the insert. RNA was purified using the Direct-zol miniprep kit (ZR2080, Zymo Research), according to the manufacturer’s specifications. HBE RNA was isolated at 4- and 7- dpi, while HNE RNA was isolated at 4- dpi along with the corresponding non-infected controls.

RNA isolated either from Vero cells (15 μg) or from epithelial culture (5 μg) was supplemented with 10ng or 3ng of bacterial RNA respectively as a spike-in control, before fragmentation using RNA Fragmentation Reagents (Thermo Fisher Scientific, AM8740). The fragmented RNA was used in RNA-seq and in m^6^A RIP-seq experiments.

Sequencing libraries for total RNA-seq/input for m^6^A RIP-seq were prepared from 10 ng of the fragmented RNA using SMARTer Stranded Total RNA-seq Kit V2, Pico Input Mammalian (Takara Bio). m^6^A RIP was performed with fragmented RNA as previously described by (Zeng et al. 2018) with m^6^A antibody (Synaptic systems, 202003) incubated with 15 μg of Vero or 5 μg of epithelial (HBE and HNE) RNA. m^6^A RIP RNA were used to make Sequencing libraries using the SMARTer Stranded Total RNA-seq Kit V2, Pico Input Mammalian (Takara Bio). All the libraries were single-end sequenced (1X 88 bp) on an Illumina NextSeq 2000 platform at the BEA core facility, Stockholm, Sweden.

### RT-qPCR and m^6^A RIP-qPCR

For performing m^6^A RIP-qPCR, 3 μg of RNA from Vero cells infected with SARS-CoV-2 B.1.1.7 variant or human patient RNA samples from throat/nose swabs (ethical permit # 2020-03276, #2020-01945 and #2022-01139-02) were used. RIP was performed with 1μg of m^6^A antibodies (synaptic systems, 202003) or IgG antibodies (SantaCruz, SC-2027). The input and m^6^A RIP-RNA were converted to cDNA using the High-Capacity RNA-to-cDNA™ Kit (Thermo Fisher Scientific, 4387406) and random primers. qPCR was performed on a Quant Studio 3 thermocycler (Thermo Fisher Scientific) using gene-specific PCR primers (**Supplemental Table S1**) mixed with Power SYBR Green Master Mix (Thermo Fisher Scientific, 4367659) and diluted cDNA (10-fold dilution) as a template. The data is represented as percentage input values.

For RT-qPCR, the RNA was directly converted to cDNA and subjected to qPCR as mentioned above, using primers listed in Supplemental Table S1. Expression values presented for each gene are normalized to *TBP* and *POL2RG* or to *GAPDH* using the delta-delta Ct method.

To check m^6^A levels in both forward and reverse strands of the viral RNA, cDNA was synthesized with primers specific for each strand separately at 50 °C (**Supplemental Table S1**) using the ImProm-II™ Reverse Transcription System (Promega, A3800) and subjected to qPCR with primers specific for the viral N-protein (**Supplemental Table S1**).

Human patient RNA samples from throat/nose swab were converted to cDNA and subjected to qPCR as mentioned above, using primers listed in Supplemental Table S1. Expression was normalized to *ACTB*.

The cellular level of SARS-CoV-2 RNA was measured by RT-qPCR in HBE and HNE infection as described previously (Samelson et al. 2022).

### G3BP1 and YTHDF2 RNA Immuno-precipitation qPCR (RIP-qPCR)

Uninfected Vero cells treated with the METTL3 inhibitor STM2457 (50μM for 48h) or with DMSO were harvested and fixed for 10 mins with formaldehyde (1% final concentration) and quenched with Glycine (0.125 M, final concentration). Fixed cells were lysed in RIPA buffer (50mM Tris pH7.4, 150mM NaCl, 0.5% Sodium Deoxycholate, 0.2% SDS, 1% IGEPAL-CA630, Protease inhibitor and RNase inhibitor) and sonicated for 10 cycles (30 secs on 30 secs off) on a Bioruptor (Diagenode) sonicator. The cleared lysate was used for RIP with anti-G3BP1 (Abcam, Ab181150), anti-YTHDF2 (Abcam, Ab246514) and IgG (SantaCruz, SC-2027) antibodies. The RNA-protein-antibody complex was captured using protein A/G magnetic beads (Thermo Fisher Scientific, 10002D and 10004D). Magnetic beads were washed with low salt buffer (1X PBS, 0.1% SDS and 0.5% IGEPAL-CA630) and high salt buffer (5X PBS, 0.1% SDS and 0.5% IGEPAL-CA630) before eluting in elution buffer (10mM Tris pH7.4, 100mM NaCl, 1mM EDTA, 0.5% SDS) with Proteinase K. RNA was extracted using TRIzol reagent and cDNA synthesis and RT-qPCR were performed as described above.

### Proximity ligation assay (PLA)

PLA was performed on Vero cells using the Duolink^®^ PLA kit (Merck, DUO92014) according to the manufacturer’s protocol, and using anti-METTL3 (Abcam, ab195352 at 1:300 dilution) and anti-METTL14 (Abcam, ab220030 at 1:500 dilution) or anti-XPO1 (Santa Cruz, sc-74454 at 1:300 dilution) antibodies. As a background control, only METTL3 or only XPO1 antibody was used in the assay. DAPI was used to mark the nuclear border and PLA foci within the nucleus were counted with ImageJ using particle analysis.

### LC-MS/MS quantification of m^6^A in viral RNA

LC-MS/MS-based quantification of m^6^A was done as previously described (Liu et al. 2020b). In brief, RNA from B.1 viral particles was isolated from Vero cell supernatant and digested by P1 nuclease (Sigma, N8630) followed by treatment with phosphatase (NEB, M0289S). The sample was then filtered (0.22 μm pore size) and directly injected into the LC-MS apparatus. As a positive standard for LC-MS/MS, and to estimate the number of m^6^A modifications in viral RNA, we parallelly processed commercially synthesized RNA oligos (**Supplemental Table S1**) with or without internal m^6^A mixed at a 5:1 A/m^6^A ratio. We made triplicate injections of the standard RNA oligos and the viral RNA and estimated the A/m^6^A ratios for both samples. LC-MS/MS profiles were monitored using the parallel reaction-monitoring (PRM) mode for: m/z 268.0–136.0, and m/z 282.0–150.1 that corresponds to A and m^6^A respectively, as previously described (Sun et al. 2021).

### METTL3 knock-down and Western blot analysis

Vero cells with stable METTL3 (METTL3-KD) and control knockdown (Control-sh) were generated using lentivirus expressing shRNA against METTL3 or non-targeting genomic regions. The shRNA sequences used in the study are provided in Supplemental Table S1.

Total proteins were extracted from cells using RIPA buffer (Sigma-Aldrich, #R0278) and quantified using the Pierce BCA Protein Assay Kit (Thermo Scientific, #23225) as per the manufacturer’s instructions. An equal amount of proteins per sample were resolved by SDS- PAGE on NuPAGE Bis-Tris gels (4-12%) (Invitrogen), followed by transfer onto 0.2 μm Nitrocellulose membrane using a Trans-Blot Turbo Transfer System (Bio-Rad). The membrane was blocked for 1 h at RT with a blocking solution (PBST-5%Milk) before incubation with the primary antibodies, anti-METTL3 (1:400, b195352; Abcam) or anti-GAPDH (1:5000, ab9485, Abcam), in blocking solution overnight at 4°C. After washing, the membranes were incubated with a secondary antibody for 1 h at RT, and the proteins were then detected with the SuperSignal West Pico PLUS Chemiluminescent Substrate (Thermo Scientific, #34579) using a ChemiDoc XRS+ system (Bio-Rad).

### Proliferation assay

5,000 cells/well were seeded in a 96-well plate to assess the proliferation of cells treated with either DMSO or Selinexor (150nM). The CellTiter 96 Non-Radioactive Cell Proliferation Assay kit (Promega, #G4000) was used to determine cell growth as per the manufacturer’s instructions. Absorbance was measured using a microplate reader Infinite 50 (Tecan, Austria).

### Processing of RNA sequencing data

Single-end sequencing reads from SMARTer-Stranded Total RNA-seq Kit v2 were analyzed with FastQC for quality control (https://www.bioinformatics.babraham.ac.uk/projects/fastqc/), and adapters were removed using Trim Galore v0.6.6 with a minimal length threshold of 20bp. Trimmed reads from Vero infected and non-infected cells were aligned to the following reference genomes: concatenated chlSab2 (Chlorosebus sabeus), wuhCor1 (SARS-CoV-2), plus Escherichia coli str. K-12 substr. MG1655 (spike-in) or the SARS-CoV-2 + E. coli genomes, obtained from the UCSC Genome Browser (http://hgdownload.soe.ucsc.edu/goldenPath/). Alignments were made using HISAT2 v2.2.1 (Kim et al. 2019) with parameters (-U --rna-strandness R). Sequencing data from human bronchial and nasal epithelial non-infected and infected cells were mapped to the concatenated human (hg38) + SARS-CoV-2 (wuhCor1) + E.coli (spike-in) reference genomes. Duplicate alignments were labeled using *markDuplicates* from *Picard* v2.23.4 and marked alignment files were further processed using Sambamba v0.7.1 (Kim et al. 2019; Tarasov et al. 2015) keeping only mapping reads separated by strand after duplicate removal.

### Analysis of RNA-seq data for differential gene expression

Aligned reads were quantified using featureCounts from the Rsubread package (Liao et al. 2019) following differential expression analysis with DESeq2 using two replicates per condition (Love et al. 2014). Genes were considered differentially expressed if they had an absolute log_2_ fold change value > 1 and an adjusted p-value cutoff of < 0.01 for Vero data and log_2_ fold change > 0.5 and adjusted p-value < 0.1 for HBE data. Heatmap visualization of differentially expressed genes was generated using the pheatmap package in R. Principal component analysis of gene expression patterns after infection with different SARS-CoV-2 variants was calculated on the regularized normalized counts obtained from DESeq2 using the factoextra package. Correlation analysis of the gene expression changes was performed between cells infected with different SARS-CoV-2 variants and also at the level of differentially expressed genes in different infection conditions. Publicly available data were used as well (Riva et al. 2020). Functional enrichment analysis was carried out using clusterProfiler (Yu et al. 2012) or Enrichr (Chen et al. 2013) against the Gene Ontology biological process or Enrichr COVID-19 related gene sets databases, respectively, with the selection of significantly enriched terms (p-value < 0.05). GO term clustering and visualization analysis was carried out using ClueGO (Bindea et al. 2013) from Cytoscape v3.8.2. Briefly, common enriched terms across any pair of variants previously identified with clusterProfiler and associated with m^6^A related genes were selected and their relationships were analyzed using ClueGO, enabling the visualization of shared genes between common terms. Calculation of network connectivity and comparison to random networks were performed using the igraph R package (https://cran.r-project.org/package=igraph). Differential exon usage analysis was performed with the DEXSeq R/Bioconductor package (Anders et al. 2012) on TPM normalized sequence counts quantified with Salmon v1.4.0 (Patro et al. 2017) in mapping-based mode using the stranded reverse library type parameter (-l SR).

### m^6^A RIP-seq data analysis

m^6^A peak calling on SARS-CoV-2 viral genome and infected cell samples was performed with callpeak from MACS2 v2.2.6 (Zhang et al. 2008) on IP and input processed alignments mapped to SARS-CoV-2 wuhCor1 or the concatenated genomes (chlSab2 + wuhCor1) or (hg38 + wuCor1), using the parameters “--no-model --keep-dup auto --call-summits” and effective genome sizes according to each genome with a p-value cutoff of 0.05 for SARS-CoV-2 genome and a p-value cutoff of 0.01 for Vero and Human datasets. The called peaks were annotated according to the nearest genomic feature using annotatePeaks.pl from HOMER v4.11 (Heinz et al. 2010). Retained, gained, and lost peaks between non-infected and infected cells were identified using BEDTools intersect from BEDTools v2.29.2 (Quinlan and Hall 2010). Briefly, peaks were classified as “retained’’ depending on whether a peak was intersected between non-infected and infected samples, “lost” if a peak was present in non-infected cells but not intersecting with peaks of infected samples, and “gained” if a peak was present in infected cells but not in non-infected cells. Motif analysis on m^6^A called peaks was carried out with findMotifsGenome.pl from HOMER and visualized with the universalmotif R package (https://bioconductor.org/packages/universalmotif/). Changes in overall m^6^A enrichment across transcripts after infection were calculated by extracting the spike-in normalized m^6^A /input signals from bigwig files within the annotated gene coordinates in the chlsab2 genomes using the rtracklayer package (Lawrence et al. 2009). Further correlation between m^6^A enrichment and gene expression changes (log_2_ fold changes infected vs non-infected) were then generated.

### Excluding potential N6,2-*O*-dimethyladenosine (m^6^Am) signals

To evaluate the presence of potential m^6^Am modification located at the first-transcribed nucleotide (Liu et al. 2020a; Tan et al. 2018), we bioinformatically excluded m^6^A RIP-seq signals located near the transcription start sites (TSS) that could correspond to m^6^Am modifications captured by the m^6^A antibody. For this, the summit coordinates of predicted m^6^A RIP-seq peaks were overlapped with the first 20 nt (close to transcription start sites) of all transcripts using the plyranges package (Lee et al. 2019) according to each reference genome (chlsab2, hg38 or wuhCor1). The m^6^A peaks located within the 20 bp regions from transcription start sites were excluded from the analysis.

### Normalization of m^6^A RIP-seq data using spike-in

To control for systematic variations across m^6^A RIP experiments, the amount of spike-in bacterial RNA was estimated by counting the total number of reads uniquely mapped to the E. coli K-12 reference genome using Sambamba v0.7.1 (Tarasov et al. 2015). E. coli spike-in counts were further used to calculate scaling factors for each batch of m^6^A RIP-seq samples (**Supplemental Table S2**). Computed scaling factors were then used in metagene density distribution analyses (described below) to normalize the density of m^6^A peaks to the spike-in content in each sample. Genome-wide coverage tracks corresponding to m^6^A IP/input ratio were further calculated and adjusted according to the computed spike-in scaling factors using bamCoverage–scaleFactor parameter from deepTools v3.3.2 (Ramírez et al. 2016).

### Quantification of viral reads

Counts of reads uniquely mapped to the SARS-CoV-2 genome were obtained with featureCounts (Liao et al. 2019) and normalized to the spike-in scaling factor. TPM values were then calculated using these counts by further normalizing with the gene lengths to obtain viral gene expression. Averaged TPM expression values were calculated for all replicates in Vero, HBE, and HNE post-infection samples, reflecting the different numbers of viral reads depending on the host cells.

### Metagene analysis

To analyze the genome-wide distribution of m^6^A, a metagene analysis of m^6^A peak density distribution was performed by overlapping the peak coordinates with the following genomic features: 5’UTR, CDS, and 3’UTR plus 1 kb upstream and downstream coordinates obtained from the GTF genome annotation files from UCSC (for chlsab2) or GENCODE v36 (for hg38), considering the longest isoform for each gene. Each transcript was scaled to fixed size metagene bins according to each reference genome. m^6^A peak density distribution profiles were generated after mapping the m^6^A peaks to the metagene coordinates using the plyranges R package (Lee et al. 2019). In order to compare multiple conditions, the relative m^6^A density distributions were calculated using the relative density function from ggmulti package (https://cran.r-project.org/web/packages/ggmulti/index.html). The relative density function calculates the sum of the density estimate area of all conditions, where the total sum is scaled to a maximum of 1 and the area of each condition is proportional to its own count. The relative m^6^A-RIP peak distributions were further normalized using the calculated spike-in factors.

### Enrichment of m^6^A peaks

To calculate the enrichment of RIP signals across m^6^A peak regions, we extracted the number of reads in m^6^A-RIP (m^6^A signal) and input RNA alignments 125 bp around peak summit coordinates using the ScoreMatrix function from the genomation package (Akalin et al. 2015). The m^6^A signal around 125 bp peak summit was normalized using spike in control to account for the differences in immunoprepitation. The obtained input RNA counts within m^6^A-peak regions were normalized to counts per million to account for differences in library sizes. The ratio of m^6^A signal/input RNA was then calculated using the enrichmentMatrix function from the same package.

### Analysis of m^6^A-readers from publicly available SARS-CoV-2 interactome

We collected publicly available data from the SARS-CoV-2 interactome in Schmidt et al., 2021 to analyze the reported interactions of known m^6^A-readers. For this, we selected proteins with a significant log_2_ enrichment over the background. As reported by the original authors, the SARS-CoV-2 RNA-protein interactome can be further divided into a core (adjusted p-value < 0.05) and expanded interactome (adjusted p-value < 0.2), containing 57 and 119 RNA interacting proteins, respectively. After matching the significantly enriched SARS-CoV-2 RNA-protein interactome data with the publicly available m^6^A-reader protein interactors collected from STRING-db, we found 16 overlapping proteins. From these, 8 proteins belong to the core SARS-CoV-2 RNA-protein interactome (YTHDF2, YBX1, SYNCRIP, PABPC1, MOV10, DDX3X, HNRNPA1, and IGF2BP2) and 8 proteins (IGF2BP1, HNRNPA3, HNRNPA0, HNRNPAB, HNRNPL, G3BP1, PCBP2, and HNRNPA2B1) form part of the expanded SARS-CoV-2 interactome. The resulting interaction network was analyzed and visualized using StringApp Cytoscape plugin (Doncheva et al. 2019).

## DATA ACCESS

The source code used for the analysis of data in this study is available in the Supplemental Code file and in the GitHub repository at: https://github.com/AkramMendez/m6a_sarscov2. All raw and processed sequencing data generated in this study have been submitted to the NCBI Gene Expression Omnibus (GEO; https://www.ncbi.nlm.nih.gov/geo/) under accession number GSE188477.

## COMPETING INTEREST STATEMENT

The authors declare no competing interests.

## ACKNOWLEDGEMENTS

We would like to thank the core facility at Novum, BEA, Bioinformatics and Expression Analysis, which is supported by the board of research at the Karolinska Institute and the research committee at the Karolinska hospital for help with sequencing; the Proteomic core facility at GU for their support with LC-MS/MS. The computations and data handling were enabled by resources in project SNIC-2022-22-85 provided by the Swedish National Infrastructure for Computing (SNIC) at UPPMAX, partially funded by the Swedish Council through grant agreement no. 2018-05973. We acknowledge Anders Sjölander for assistance concerning technical and implementational aspects of the UPPMAX resources.

## FUNDING

This work was funded by grants from the Swedish Research Council (Vetenskapsrådet) to TM (2018-02224), Sweden-South Korea COVID-19 grant from the Swedish Research Council (Vetenskapsrådet) to TM (2020-06311, and 2022-05965) and project grant to TM from Svenska Läkaresällskapet; Kungl. Vetenskaps-och Vitterhets-Samhället (KVVS) grant to TM, grants COROCHIP and PFR7 from the Urgence COVID-19 Fundraising Campaign of Institut Pasteur to L.A.C; Bollan scholarship to RV.

## Notes

https://www.ncbi.nlm.nih.gov/geo/GSE188477

### Competing Interest Statement

The authors have declared no competing interest.

