## Supplemental Material for "Global loss of cellular m^6^A RNA methylation following infection with different SARS-CoV-2 variants"

Vaid, Mendez, *et al*

Department of Clinical Chemistry and Transfusion Medicine, Sahlgrenska University Hospital,

Gothenburg University, Gothenburg, Sweden.

#### Contents

Supplemental Figures S1-S7

Supplemental Table S1: Sequence of oligos used in the study.

Supplemental Table S2: Scaling factors used for normalizing m$^6$A RIP-seq data.

#### Other Supplementary Materials for this manuscript

Supplemental Data S1: Differentially expressed genes following SARS-CoV-2 infection in Vero and HBE cells

Supplemental Data S2: Enrichment analysis of differentially expressed genes following SARS-CoV-2 infection in Vero and HBE cells.

Supplemental Data S3: m$^6$A peaks in the SARS-CoV-2 viral genome and in the host genomes of non-infected and SARS-CoV-2 infected Vero, HBE and HNE.

Supplemental Data S4: Genes with differential exon usage (DEU) in SARS-CoV-2 infected Vero cells.

#### Supplemental Figure S1

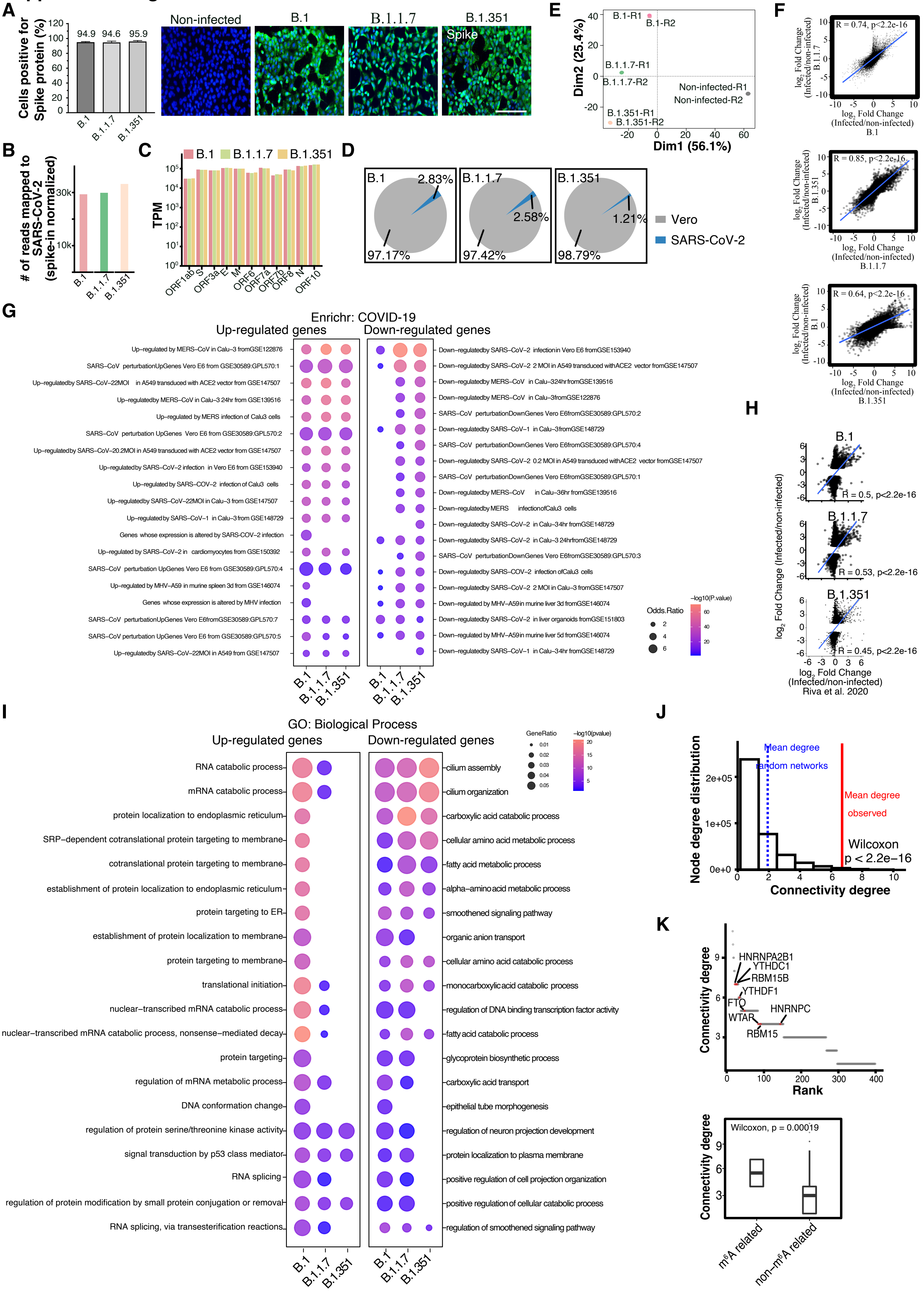

**Supplemental_Figure_S1** (A) SARS-CoV-2 Spike protein staining summarized as bar plots with mean ± SD shown (left panel; n=2) and corresponding representative images of Spike staining (right panel) in Vero cells 48 h post-infection with three different SARS-CoV-2 variants as indicated. The scale bar is 200 μm. (B) Bar plots showing spike-in normalized reads mapped to the SARS-CoV-2 genome in Vero cells infected with different variants of SARS- CoV-2. (C) The $\log_{10}$ transcripts per million (TPM) values for SARS-CoV-2 genes in B.1, B.1.1.7, and B.1.351 infected cells. Data shown as the mean of two replicates. (D) Pie chart showing the percentage of reads mapping to host (Vero) and viral (SARS-CoV-2) genomes. (E) Principal-component analysis (PCA) of global expression patterns of SARS-CoV-2 infected (variants as specified) and non-infected Vero cells. Replicates are labelled with R1 and R2 suffixes. (F) Scatter plots showing correlation between $\log_2$ fold changes in gene expression after infection with different SARS-CoV-2 variants in Vero cells. Statistics: Pearson's correlation test. The blue line shows the linear regression with 95% confidence interval. (G) Top enriched terms associated with up-regulated and down-regulated genes after SARS-CoV-2 infection, based on publicly available COVID-19 related gene sets. Data were obtained from the Enrichr database. (H) Scatter plots of differentially expressed genes depicting the correlation between the $\log_2$ fold changes in gene expression after B.1, B.1.1.7, and B.1.351 infection and the $\log_2$ fold changes reported in publicly available data of SARS- CoV-2 infected Vero cells 24-hour post infection (Riva et al., 2020). The blue line depicts the linear regression line with 95% confidence interval. Statistics: Pearson's correlation test. (I) Top GO biological process terms associated with up-regulated and down-regulated genes after infection of Vero cells with the B.1, B.1.1.7, and B.1.351 strains. The size of the dots represents the enrichment of genes for a given GO term; the dots are colored according to their significance in $\log_{10}$ p-value. (J) Node connectivity degree distribution of 1000 randomly generated networks of the same size. The mean connectivity of random networks (blue dotted line) was compared to the observed mean connectivity degree of the network obtained after ClueGO analysis for RNA catabolism-associated genes (red line). Statistical significance was calculated using the Wilcoxon signed rank test. (K, top panel) Ranking of nodes according to their connectivity degree, with $m^6A$-related proteins highlighted in red. (K, bottom panel) Connectivity of $m^6A$-related proteins compared to other proteins in the network. Statistical significance was calculated using the Wilcoxon test.

### Supplemental Figure S2

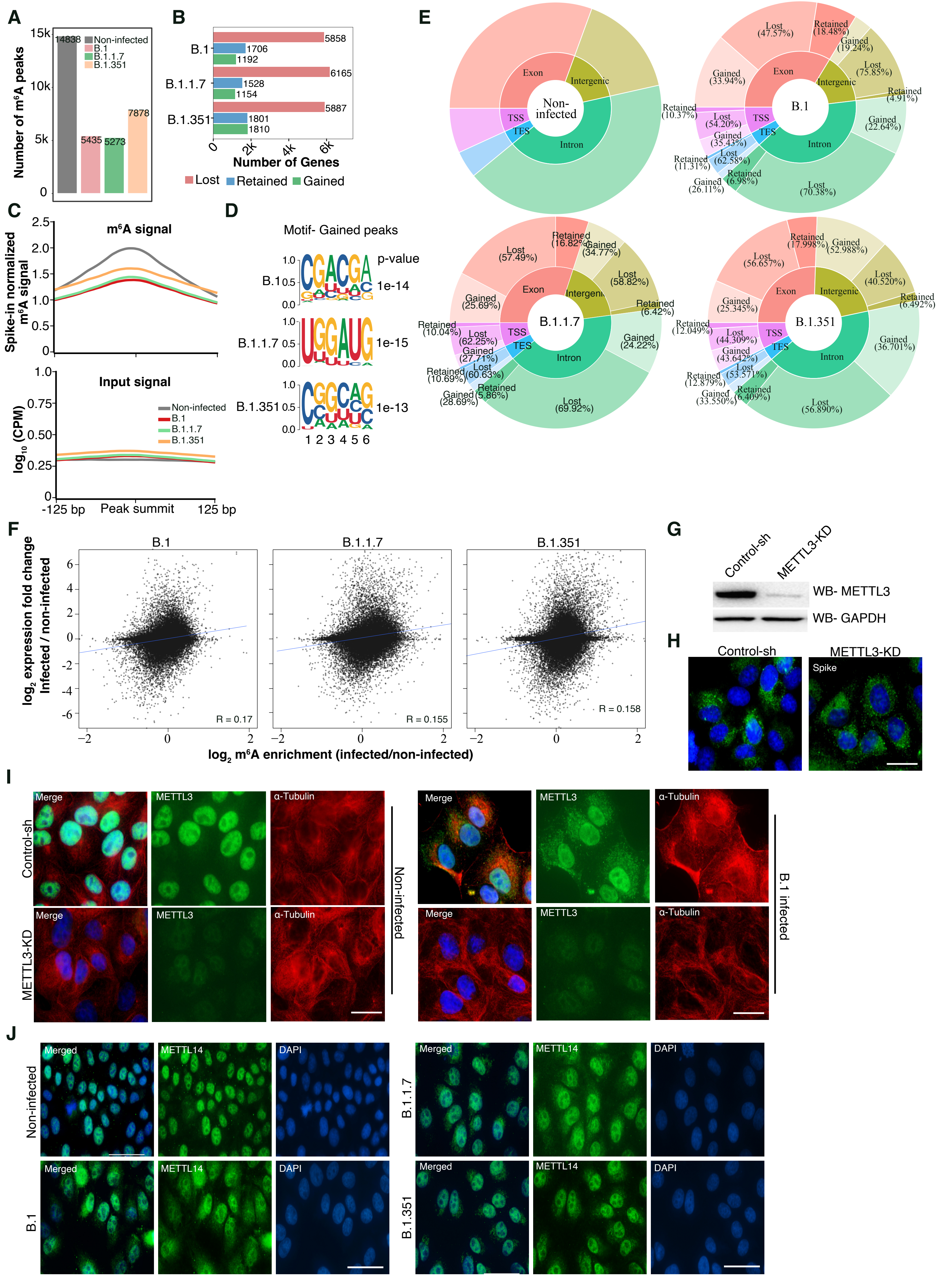

**Supplemental_Figure_S2** (A) Number of $m^6A$ peaks in non-infected and infected Vero cells. (B) Bar plots representing the number of genes with lost, gained, or retained $m^6A$ peaks in Vero cells after infection with three different SARS-CoV-2 variants, as indicated. (C)S p i k e - i n normalized $m^6A$ signal (top panel) and corresponding input signal ($log_{10}$ CPM) (bottom panel) at the $m^6A$ peak regions (± 125bp from $m^6A$ peak summit) that were commonly lost post-infection with the three different variants of SARS-CoV-2 compared to non-infected Vero cells. (D) Identified motifs from *de novo* motif analysis of gained $m^6A$ peaks. Motifs from Vero cells infected with B.1, B.1.1.7, and B.1.351 are shown. (E) Donut plot distribution of annotated $m^6A$ peaks by genomic features for non-infected and infected Vero cells with three different variants, as indicated. The proportion of retained, gained, and lost $m^6A$ peaks annotated in exonic, intronic, transcription start site (TSS), transcription end site (TES), and intergenic regions are shown. (F) Scatter plots depicting the correlation between $log_2$ fold changes in gene expression and $log_2$ fold changes in $m^6A$ enrichment after infection with B.1, B.1.1.7, and B.1.351 variants. The blue line depicts linear regression line with 95% confidence interval. Statistics: Pearson's correlation test. (G) Western-blot validation of METTL3 knock-down in Vero cells, with GAPDH used as a loading control. (H) Immunostaining using Spike antibody in control and METTL3-KD cells 48 h post SARS-CoV-2 infection. The scale bar is 20 µm (I) Immunostaining for METTL3 and α-Tubulin in Control and METTL3-KD Vero cells that were either non-infected or infected with the B.1 SARS-CoV-2 variant. The scale bar is 20 µm (J) Immunostaining showing METTL14 localization in non-infected and Vero cells infected with the B.1, B.1.1.7, B.1.351 variants at 24h. The scale bar is 50 µm.

### Supplemental Figure S3

**A**

| Sample | Expected ratio A/m6A | Transition A | Transition m6A | Ratio A/m6A | Average ratio | Avg. number of m6A in 8954 Adenine in viral genome |
|---|---|---|---|---|---|---|
| Standard 1 | 5 | 2.26e+05 | 4.38e+04 | 5.2 | | |
| Standard 2 | 5 | 2.30e+05 | 3.59e+04 | 6.4 | 5.4 | N/A |
| Standard 3 | 5 | 2.32e+05 | 4.87e+04 | 4.8 | | |
| SARS-CoV-2 B.1 Exp1 | N/A | 4.26e+07 | 6.09e+04 | 698.7 | | |
| SARS-CoV-2 B.1 Exp2 | N/A | 4.45e+07 | 4.41e+04 | 1,008.0 | 832.0 | 10.75* +/-1.98 |
| SARS-CoV-2 B.1 Exp3 | N/A | 3.87e+07 | 4.89e+04 | 790.3 | | |

**B**

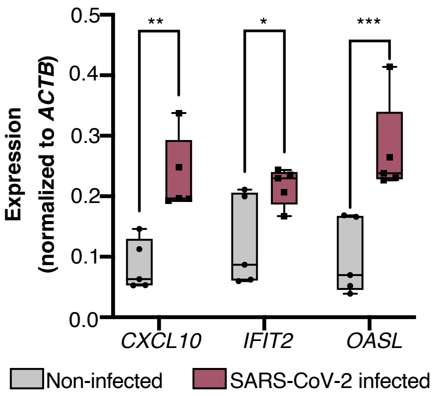

**C**

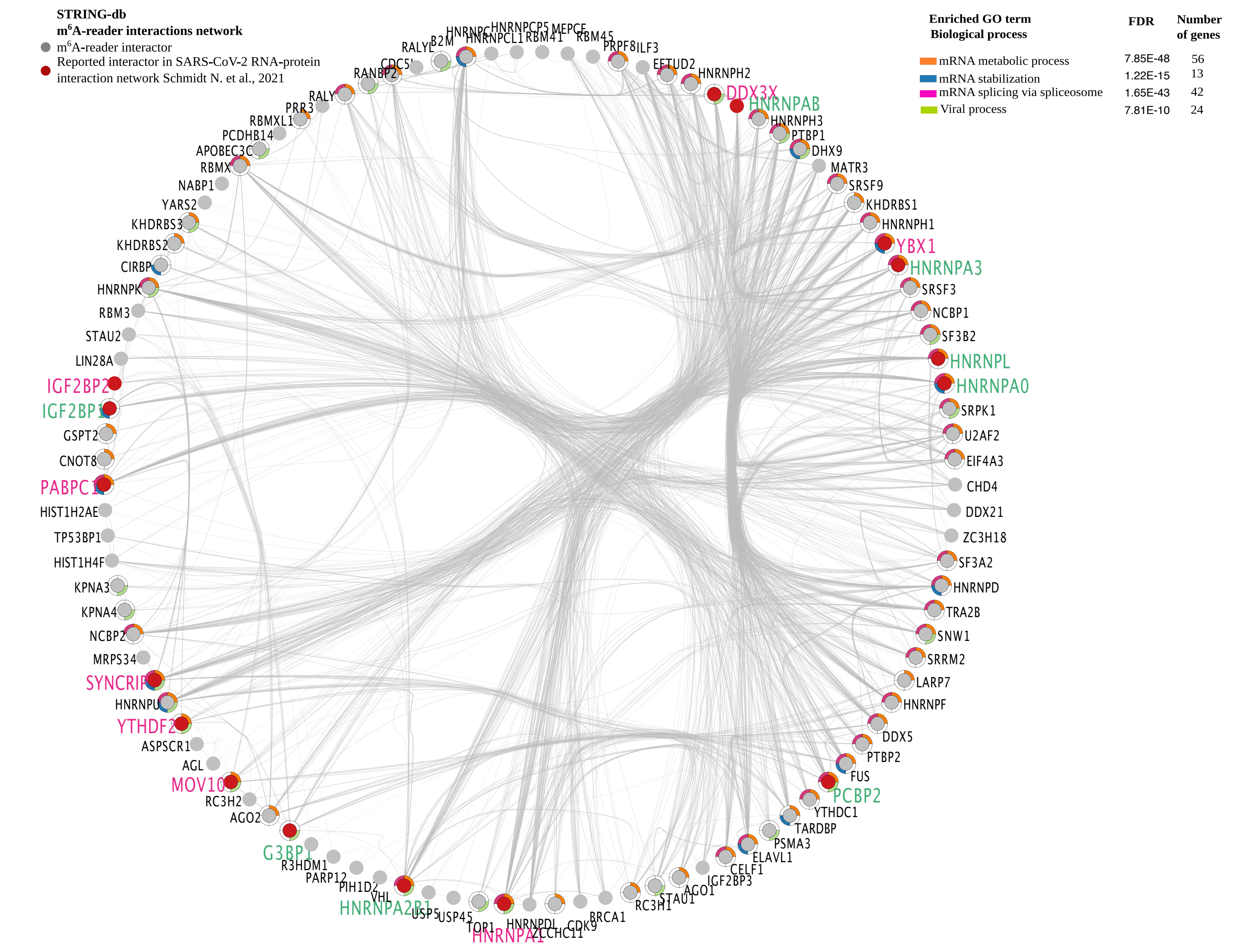

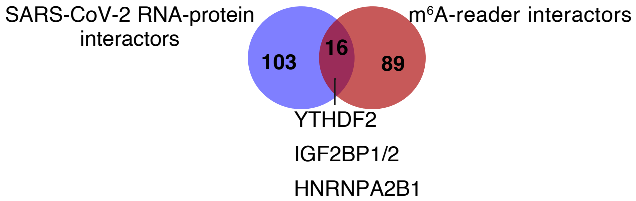

**Supplemental_Figure_S3** (A) Summary of LC-MS/MS quantification of $m^6A$ in control standard RNA oligos and B.1 viral genomic RNA. The average ratio of A/ $m^6A$ in the viral genomic RNA was 832. Considering the total number of 'A' in the viral genome to be 8954, the number of 'A' with $m^6A$ modification is on average 10.75 per viral genome. (B) RT-qPCR expression data (normalized to *ACTB*) of three selected genes in RNA isolated from throat/nose swab samples who were either positive (SARS-CoV-2 infected) or negative (Non-infected) for COVID-19. (C) Interaction network of $m^6A$-related proteins. (Top panel) Interaction network of $m^6A$-related proteins. Known $m^6A$-reader interacting proteins collected from the STRING- db (grey nodes) were matched with the SARS-CoV-2 RNA-protein interactome obtained from Schmidt et al., 2020 (red nodes). The $m^6A$-related proteins pertaining to the core or the expanded SARS-CoV-2 interactome (as reported by Schmidt et al., 2021) labeled in pink and green, respectively. Outer node colors indicate enriched GO biological process terms for the $m^6A$-related proteins of the network. (Bottom panel) The Venn diagram shows the number of overlapping proteins between two datasets as indicated.

### Supplemental Figure S4

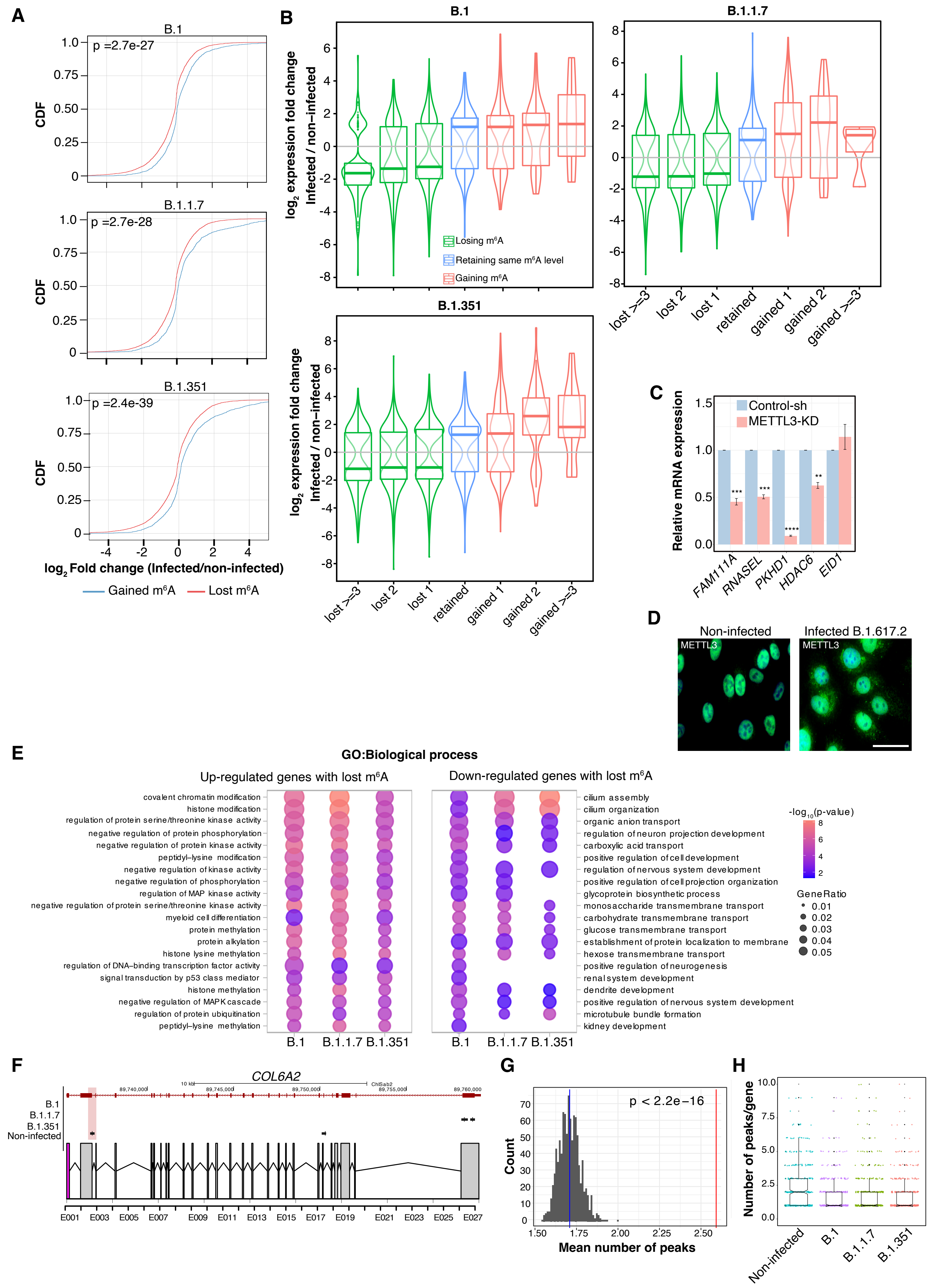

**Supplemental_Figure_S4** (A) Cumulative distribution function (CDF) plots showing the cumulative density distribution of $\log_2$ fold changes in gene expression, with a comparison of lost (red) and gained (blue) $m^6A$ genes after infection with different SARS-CoV-2 variants. Statistical significance was calculated using the Wilcoxon test. (B) Expression of $m^6A$-modified genes after SARS-CoV-2 infection. The distribution of differentially expressed genes categorized according to changes in $m^6A$ levels after viral infection compared to non-infected cells is shown. Genes were categorized as gaining one or more peaks (non- $m^6A$ genes in non-infected cells being $m^6A$ -modified after infection), losing one or more peaks ($m^6A$ -modified genes in non-infected cells showing a decrease in $m^6A$ peak number), or retaining $m^6A$ (showing the same number of $m^6A$ peaks in non-infected and infected cells). (C) Relative expression of selected genes with varying levels of $m^6A$ modification in METTL3-KD Vero cells compared to control cells. Data was normalized to *GAPDH* expression. Statistics: two- tailed *t*-test; **: $p < 0.01$, ***: $p < 0.001$, ****: $p < 0.0001$, n=3. (D) METTL3 localization in Vero cells infected with the B.1.617.2 (Delta) variant. The scale bar is 50 µm. (E) Top enriched terms associated with up-regulated and down-regulated genes with lost $m^6A$ upon infection. The size of the dots depicts the gene enrichment ratio, while their color indicates the $\log_{10}$ level of significance. (F) Differential exon usage for the *COL6A2* gene upon SARS-CoV-2 infection. The genome browser visualization (top) shows the localization of $m^6A$ peaks in non-infected and infected cells along the *COL6A2* gene. The exon structure (bottom) with a predicted differentially used exon (DEU) (in pink) included after infection is shown. (G) Distribution of the number of $m^6A$ peaks over DEU genes compared to a random distribution in $m^6A$ positive genes (1000 replicates without replacement). The average number of $m^6A$ peaks in DEU genes and in $m^6A$ positive random genes are denoted by a red and blue line, respectively. Statistics: one sample *t*-test. (H) Distribution of $m^6A$ peak numbers in DEU genes in non-infected and infected Vero cells (infection with different variants as indicated).

### Supplemental Figure S5

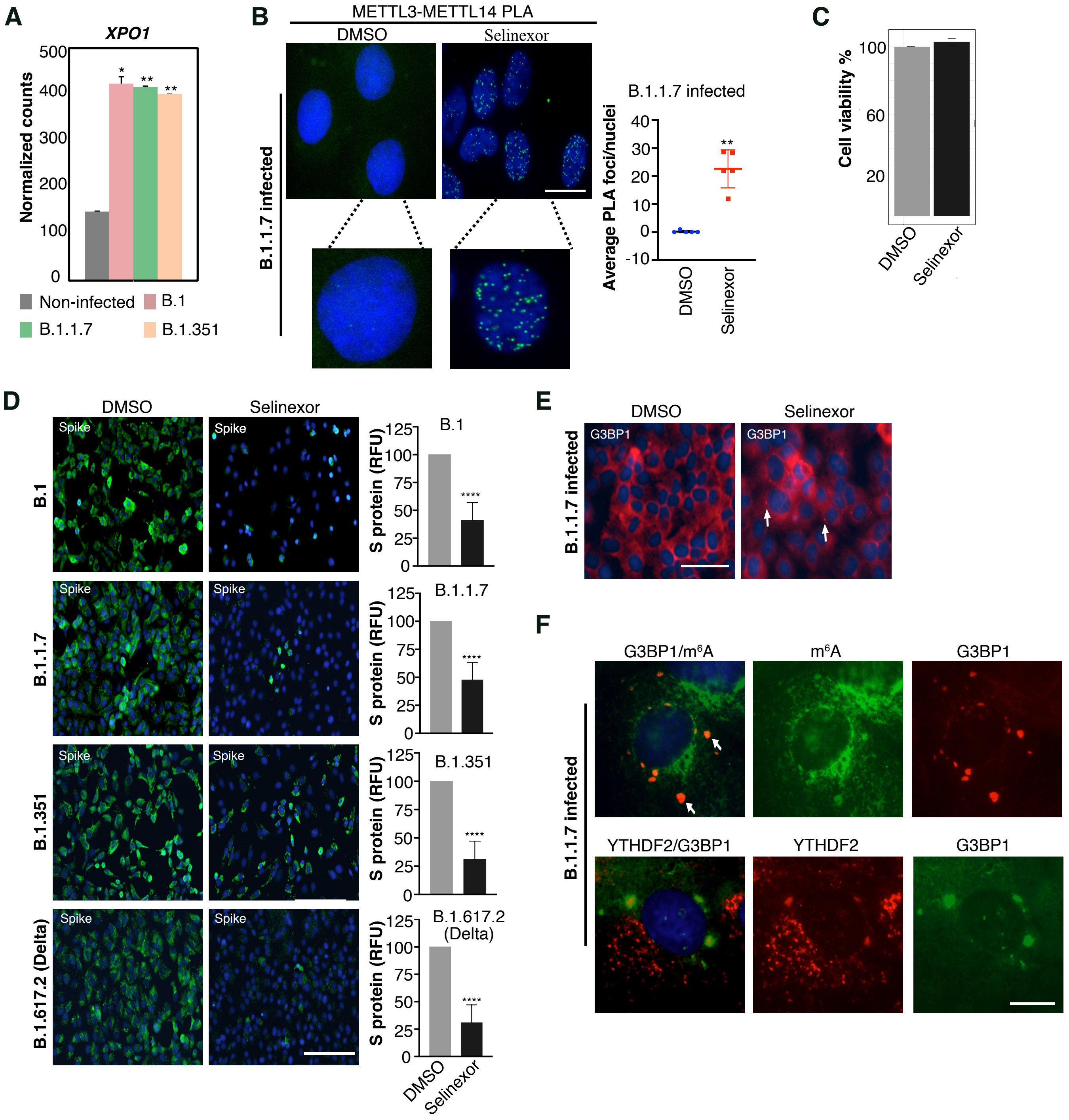

**Supplemental_Figure_S5** (A) Bar graph depicting the expression of *XPO1* after infection with the B.1, B.1.1.7, and B.1.351 variants. Normalized read counts in RNA-seq data are reported. (B, left panel) PLA experiment revealing METTL3 and METTL14 interaction foci in the nucleus (marked by DAPI) of Vero cells infected with the B.1.1.7 variant and treated with either DMSO or Selinexor (150nM). The scale bar is 20 µm. (B, right panel) Quantification of METTL3-METTL14 PLA foci as detected in the left panel. The number of PLA foci/nuclei are shown as Mean ± SD. Data presented from multiple experiments with the total number of cells counted ≥100. Statistics: one-way ANOVA, **: $p < 0.01$. (C) MTT assay showing Vero cell viability after Selinexor treatment compared to DMSO control. (D, left) Immunostaining showing SARS-CoV-2 Spike protein in Vero cells at 48 h post-infection with the B.1, B.1.1.7, and B.1.617.2 (Delta) variants, with or without Selinexor treatment. (D, right) Quantification of Spike protein fluorescence in Selinexor-treated and untreated cells infected with different variants, as indicated. Spike fluorescence was quantified in 5 different fields obtained from several experiments. The scale bar is 200 µm. (E) Immunostaining showing G3BP1 localization after Selinexor treatment in B.1.1.7-infected cells at 24 h post-infection. White arrows highlight some of the G3BP1 foci. The scale bar is 50 µm. (F) Immunostaining showing the localization of G3BP1 and $m^6A$ or YTHDF2 after DMSO or Selinexor treatment in B.1.1.7-infected cells at 24 h post-infection. White arrows highlight some of the G3BP1 foci overlapping with $m^6A$ signal. The scale bar is 10 µm.

#### Supplemental Figure S6

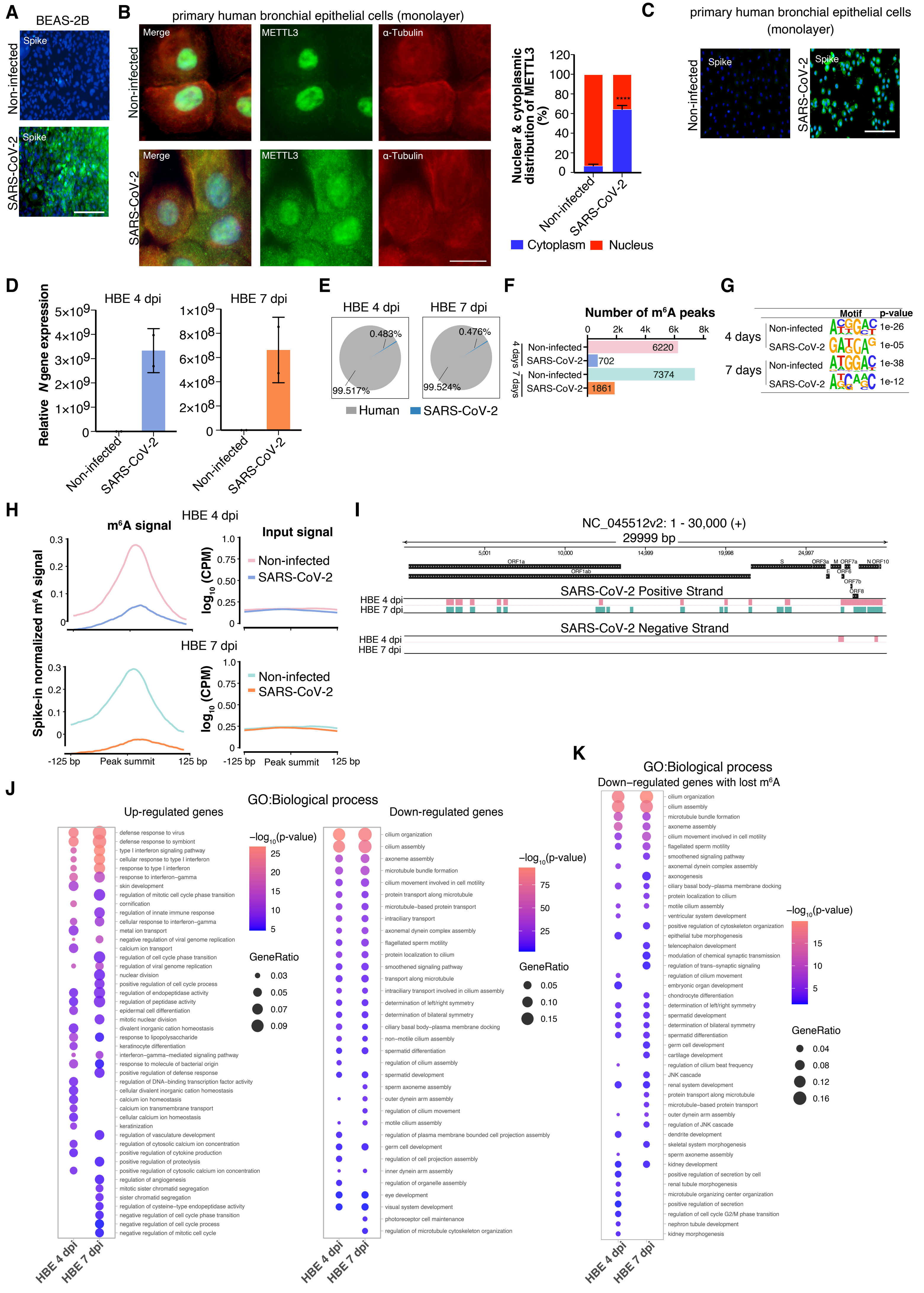

**Supplemental_Figure_S6** (A) Spike immunostaining in BEAS-2B cells that were infected with SARS-CoV-2. The scale bar is 200 µm. (B, left panel) METTL3 localization in monolayer culture of non-infected human primary bronchial epithelial cells and after infection with SARS-CoV-2. The scale bar is 20 µm. (B, right panel) The percentage distribution of METTL3 labeling in the nucleus and cytoplasm was calculated with ImageJ, using DAPI as a nucleus marker and α-Tubulin as a cytoplasm marker. Data are shown as mean ± SD. Data were obtained from multiple experiments with the total number of cells counted ≥110. Statistics: unpaired *t*-test; ****: $p < 0.0001$. (C) Spike protein immunostaining in human primary bronchial epithelial cells grown in monolayer that were infected with SARS-CoV-2. The scale bar is 200 µm. (D) Relative expression of SARS-CoV-2 *N* gene in non-infected and SARS- CoV-2 infected HBE samples 4- and 7- dpi. (E) Pie chart showing the percentage of RNA-seq reads mapping to host (Human) and viral (SARS-CoV-2) genomes in infected HBE samples. (F) Bar plot depicting the number of $m^6A$ peaks in non-infected and SARS-CoV-2 infected HBE at 4- and 7- dpi. (G) Motif analysis of the $m^6A$ peaks in non-infected and infected HBE. (H) Spike-in normalized $m^6A$ signal (left panel) and the corresponding input signal ($\log_{10}$ CPM) (right panel) at the $m^6A$ peak regions (± 125bp from $m^6A$ peak summit) that were lost in infected HBE at 4- dpi (top) and 7- dpi (bottom) compared to non-infected HBE. (I) Presence of $m^6A$ in the positive and negative strands of the SARS-CoV-2 viral genome at 4- and 7- dpi HBE cells. $m^6A$ peak regions are indicated as colored rectangles. (J) Top GO biological process terms associated with up-regulated and down-regulated genes in HBE at day 4- and 7- dpi with SARS-CoV-2. (K) Top enriched terms associated with lost $m^6A$ genes that were down- regulated in HBE at day 4- and 7- dpi with SARS-CoV-2. (J, K) The size of the dots depicts the gene enrichment ratio, while the color of the dots indicates the $\log_{10}$ level of significance.

#### Supplemental Figure S7

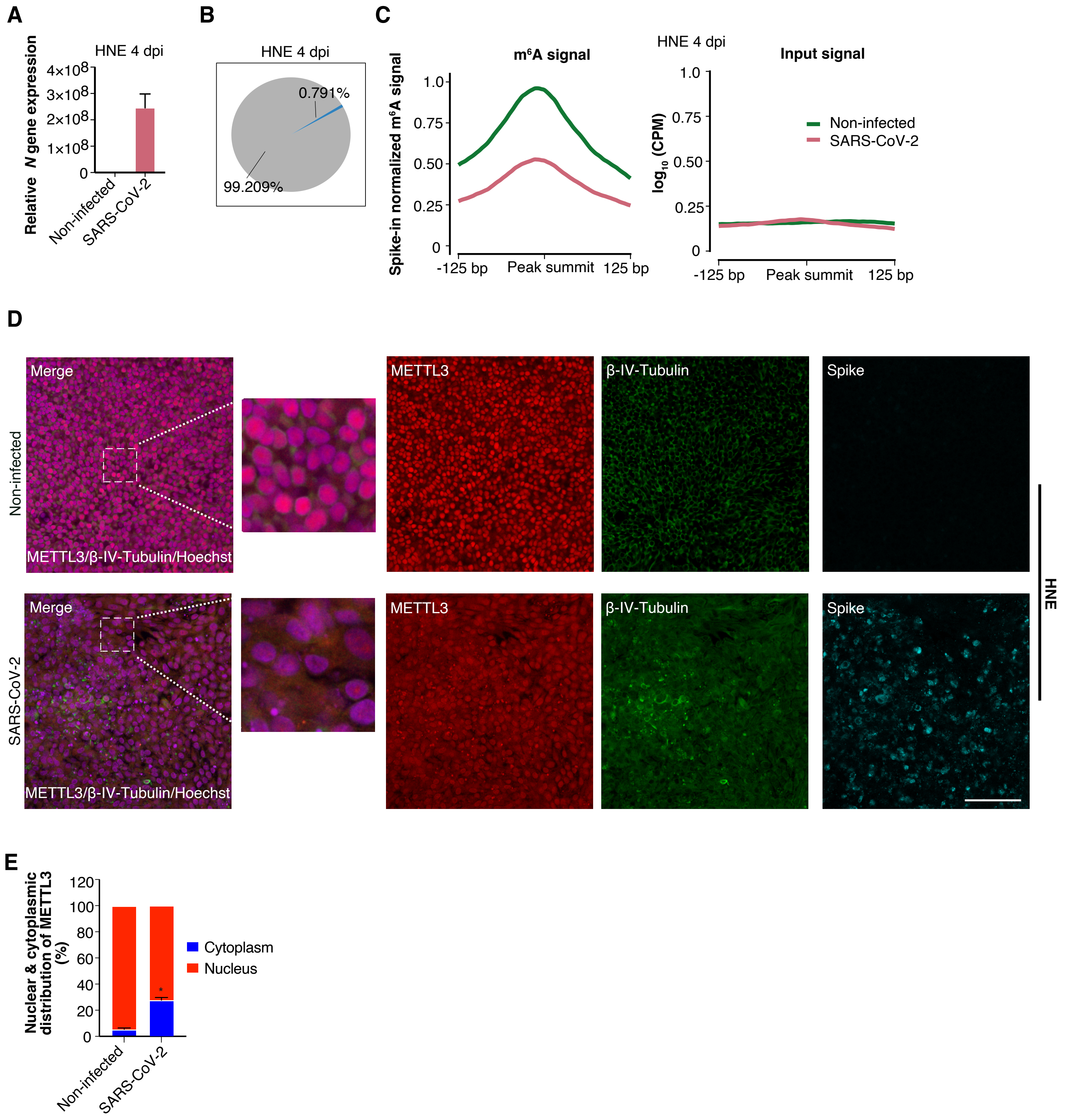

**Supplemental_Figure_S7** (A) Relative expression of SARS-CoV-2 *N* gene in non-infected and SARS-CoV-2 infected HNE samples at 4- dpi. (B) Pie chart showing the percentage of reads mapping to the host (Human) and viral (SARS-CoV-2) genomes in infected HNE at 4- dpi. (C) Spike-in normalized m$^6$A signal (left panel) and the corresponding input signal ($log_{10}$ CPM) (right panel) at the m$^6$A peak regions (± 125bp from m$^6$A peak summit) that were lost at 4- dpi in infected compared to non-infected HNE. (D) Immunostaining staining showing METTL3 (in red), β-IV-Tubulin (in green) and SARS-CoV-2 spike protein (in cyan) in HNE 4- dpi along with non-infected control. Merged image represents METTL3 and β-IV-Tubulin along with Hoechst. The scale bar is 200 μm. (E) The percentage distribution, of METTL3 in the nucleus and cytoplasm, was calculated using ImageJ. Hoechst was used as a nuclear marker and β-IV-Tubulin marking the cytoplasm. Data are shown as mean ± SD. Data were acquired from n=2 experiments with the total number of cells counted ≥ 1000. Statistics: unpaired t-test; ****: $p < 0.0001$.

#### Supplementary Tables

Supplementary Table S1: Sequence of oligos used in the study.

| Vero | | |
|---|---|---|
| **Primer name** | **Sequence** | |
| *FAM111A*-Forward primer | CCTTTCCTTCTGGCTCTTCA | |
| *FAM111A*-Reverse primer | TGTGGGAGACGGAATAGAGC | |
| *HDAC6*-Forward primer | CCCAATCTAGCGGAGGTAAA | |
| *HDAC6*-Reverse primer | CGCGATTAGGTCTTCTTCCA | |
| *PKHD1*-Forward primer | GTGGGCATTTGGTCTGAAAG | |
| *PKHD1*-Reverse primer | AGTTGTCCCAGCAGGACAGT | |
| *RNASEL*-Forward primer | TGTCAATGTGAGGGGAGAAA | |
| *RNASEL*-Reverse primer | CTTCACCAAACCCAAGTGCT | |
| *EID1*-Forward primer | TCGCCTCCTTTTTCACAACT | |
| *EID1*-Reverse primer | TGCCATTGAAAAACTTGACCT | |
| *XPO1*-Forward primer | TGGTACAGAATGGTCATGGAA | |
| *XPO1*-Reverse primer | TCTTCATGCATTGCTCCACT | |
| *YTHDF1*-Forward primer | TGGGACAAATGTGAACATGC | |
| *YTHDF1*-Reverse primer | TTTAGGCTGTGGTTTTGCAG | |
| *FTO*-Forward primer | GGAACCTTATTTTGGCATGG | |
| *FTO*-Reverse primer | GCTGACCTGTCCACCAGATT | |
| *SPEN*-Forward primer | CCACTCTCTCGGGATCAAAA | |
| *SPEN*-Reverse primer | AATCTCGTTTGGAGCGCTAT | |
| *GAPDH*-Forward primer | ATGTTCGTCATGGGTGTGAA | |
| *GAPDH*-Reverse primer | GGTGCTAAGCAGTTGGTGGT | |
| *POL2RG*- Forward primer | ACTGTTGGTGGGTGAGCAC | |
| *POL2RG*- Reverse primer | CCAGACTGGCAGCAAGAAAA | |
| *TBP*- Forward primer | ACTGTTGGTGGGTGAGCAC | |
| *TBP*- Reverse primer | CCAGACTGGCAGCAAGAAAA | |
| SARS-Cov-2 | | |
| **Primer name** | **Sequence** | |
| *N*-protein-Forward primer | CACATTGGCACCCGCAATC | |
| *N*-protein-Reverse primer | GAGGAACGAGAAGAGGCTTG | |
| *N*-Forward strand | CAGCACTGCTCATGGATTG | Primer for cDNA synthesis |
| *N*-Reverse strand | GACCCCAAAATCAGCGAAAT | Primer for cDNA synthesis |
| Human | | |
| **Primer name** | **Sequence** | |
| *CXCL10* Forward primer | TGATCTCAACACGTGGACAA | |

| | |
|---|---|
| *CXCL10* Reverse primer | ACTGTACGCTGTACCTGCAT |
| *IFIT2* Forward primer | AAAGGAACCAGAGGCCACTT |
| *IFIT2* Reverse primer | GCTCGGTTCAGGCAGCTG |
| *OASL* Forward primer | GGACTCTCTGCTCCATCCTC |
| *OASL* Reverse primer | GCAGCCAAGCATCACAAAGA |
| *ACTB* Forward primer | CTCGTAGCTCTTCTCCAGGG |
| *ACTB* Reverse primer | GGGAAATCGTGCGTGACATT |
| RNA standard for LC-MS/MS | |
| RNA oligo with $m^6A$ | GG$m^6$ACUAA$m^6$ACU |
| RNA oligo without $m^6A$ | GGACUAAACU |
| shRNA sequence | |
| *METTL3* shRNA | GCTGCACTTCAGACGAATTAT |
| Control shRNA | ATCTCGCTTGGGCGAGAGTAAG |

Supplementary Table S2: Scaling factor used for normalizing m$^6$A RIP data

| Vero | |
|---|---|
| **Sample** | **Scaling factor** |
| Non-infected | 1 |
| B.1 | 0,72 |
| B.1.7.7 | 0,81 |
| B.1.351 | 0,98 |
| HBE | |
| **Sample** | **Scaling factor** |
| HBE Non-infected day 4 | 0,75 |
| HBE SARS-CoV-2 day 4 | 1 |
| HBE Non-infected day 7 | 0,99 |
| HBE SARS-CoV-2 day 7 | 1 |
| HNE | |
| **Sample** | **Scaling factor** |
| HNE Non-infected | 0,92 |
| HNE SARS-CoV-2 day 4 | 1 |

**Supplemental Data**

**Supplemental Data S1: Differentially expressed genes**

All the differentially expressed genes post SARS-CoV-2 infection in Vero cells and HBE.

**Supplemental Data S2: Enrichment analysis**

Enrichment analysis of differentially expressed genes following SARS-CoV-2 infection in Vero and HBE cells.

**Supplemental Data S3: $m^6A$ peaks**

$m^6A$ peaks identified in SARS-CoV-2 (viral genome) and in non-infected and SARS-CoV-2 infected Vero and HBE cells (host genomes).

**Supplemental Data S4: DEU**

Genes with differential exon usage (DEU) in SARS-CoV-2 infected cells.
